# TRAIL-induced cytokine production via NFKB2 pathway promotes neutrophil chemotaxis and immune suppression in triple negative breast cancers

**DOI:** 10.1101/2024.07.19.604341

**Authors:** Manjari Kundu, Yoshimi E. Greer, Alexei Lobanov, Lisa Ridnour, Renee N. Donahue, Yeap Ng, Shashi Ratnayake, Donna Voeller, Sarah Weltz, Qingrong Chen, Stephen J. Lockett, Maggie Cam, Daoud Meerzaman, David A. Wink, Roberto Weigert, Stanley Lipkowitz

## Abstract

Tumor necrosis factor-related apoptosis-inducing ligand (TRAIL) is a potential cancer therapeutic that induces apoptosis in cancer cells while sparing the non-malignant cells in preclinical models. However, its efficacy in clinical trials has been limited, suggesting unknown modulatory mechanisms responsible for the lack of TRAIL activity in patients. Here, we hypothesized that TRAIL treatment elicits transcriptional changes in triple negative breast cancer (TNBC) cells that alter the immune milieu. To test this, we performed an RNAseq analysis of MDA-MB-231 cells treated with TRAIL, followed by validation in additional TNBC cell lines. TRAIL significantly induces expression of multiple cytokines such as CXCLs 1, 2, 3, 8,11 and IL-6, which are known to modify neutrophil function. Mechanistically, the induction of these cytokines was predominantly mediated by death receptor 5, caspase 8 (but not caspase 8 enzymatic activity), and the non-canonical NFKB2 pathway. The cytokines produced by the TRAIL-treated TNBC cells enhanced chemotaxis of healthy human donor isolated neutrophils. *In vivo*, TRAIL treated TNBC murine xenograft tumors showed activation of the NFKB2 pathway, elevated production of CXCLs and IL-6, and increased neutrophil recruitment into the tumors. Moreover, donor isolated neutrophils preincubated in supernatants from TRAIL-treated TNBC cells exhibited impaired cytotoxic effect against TNBC cells. Transcriptomic analysis of neutrophils incubated with either TRAIL alone or supernatant of TRAIL-treated TNBC cells revealed increased expression of inflammatory cytokines, immune modulatory genes, immune checkpoint genes, and genes implicated in delayed neutrophil apoptosis. Functional studies with these neutrophils confirmed their suppressive effect on T cell proliferation and an increase in Treg suppressive phenotype. Collectively, our study demonstrates a novel role of TRAIL-induced NFKB2-dependent cytokine production that promotes neutrophil chemotaxis and immune suppression.

## Introduction

Triple-negative breast cancer (TNBC), defined by the absence of estrogen receptor, progesterone receptor and HER2 amplification, is associated with a worse prognosis due to its frequent metastatic relapse after treatment of early stage disease and limited effective therapies in the advanced stage [1]. New targeted therapies including trophoblast antigen 2 directed antibody drug conjugates and immune check point inhibitors have significantly improved the outcomes of patients with TNBC [2–6]. However, identifying additional novel therapies for advanced TNBC remains an unmet need.

Tumor necrosis factor-related apoptosis-inducing ligand (TRAIL/Apo2L/TNFSF10) is a member of the TNF family of ligands which triggers caspase-mediated apoptosis through Death Receptor 4 (DR4/TRAILR-1/TNFRSF10A) and Death Receptor 5 (DR5/TRAILR- 2/TNFRSF10B) [7, 8]. TRAIL induces apoptosis in diverse cancer cells while sparing the normal cells [7, 9–12]. Previous studies showed that TNBC with a mesenchymal phenotype (a.k.a, Basal B TNBC) are the most sensitive to TRAIL-induced apoptosis [13–17]. Despite strong preclinical data, TRAIL agonists alone or in combination with chemotherapy have not shown significant activity in clinical trials in many cancers, including TNBC [18, 19]. This raises the critical question about the mechanisms that prevent TRAIL activity in human tumors.

Apart from the direct anti-tumor apoptotic effect, several studies have demonstrated that TRAIL has both anti-tumorigenic and pro-tumorigenic effects on the tumor microenvironment mediated by immune modulation, thus acting as a double-edged sword [20–22]. Increases in inflammatory cytokines such as IL8, TNFA, MIP-2, MIP-1B have been reported by activation of TRAIL-receptors in different cancer cell lines in an NFKB-dependent manner [23]. The endogenous TRAIL/TRAIL-R system was reported to produce CCL2 cytokine promoting recruitment of immune suppressive MDSCs and M2 polarized monocytes [24]. Recently, high dose recombinant murine TRAIL was reported to promote cytokine production by tumors and recruitment of pro- tumorigenic M2-like macrophage [25].

Neutrophils, the most abundant leukocytes in the blood [26], are a significant component of the tumor microenvironment, and play a pivotal role in breast cancer targeted therapies [27, 28]. The presence of tumor associated neutrophils (TANs), TAN related gene expression, and a high neutrophil to lymphocyte ratio are associated with immune suppressive activity and poor outcomes in breast cancer patients [29–31]. Mouse models of TNBC demonstrated a significant increase in neutrophil extracellular traps in peripheral blood and lung tissue augmenting the metastatic potential of the TNBC cells[32]. However, few studies report detailed insights on the mechanistic and functional significance of TRAIL induced neutrophil modulation in TNBC.

Here we demonstrated a novel role of TRAIL-induced NFKB2 dependent cytokine production by TNBC cells, promoting neutrophil recruitment. We also report the direct effect of TRAIL on the neutrophil transcriptome potentially causing immune suppression. Hence, we propose that alterations in the innate immune system may modulate the effects of TRAIL on TNBC tumors.

## Results

### TRAIL induces transcriptomic change in TNBC cell lines

To decipher TRAIL induced transcriptional changes in TNBC cells, we performed a time course of RNA expression in the TRAIL-sensitive TNBC cell line, MDA-MB-231 treated with TRAIL (45 nM; a sub-IC50 dose to observe the differential gene expression in live TRAIL-treated cells instead of apoptotic cells (Fig. 1A, upper panel and Supplementary Fig. 1A). TRAIL-dependent transcriptomic changes in MDA-MB-231 were determined at 0, 1h, 3h, 6h, 12h and 24h. Our analysis as illustrated by Heat map revealed that incubation with TRAIL significantly altered gene expression at 3h, 6h, 12h and 24h compared to 0h, whereas there was no significant difference in gene expression between 0h and 1h (Fig. 1A, lower panel, Supplementary Fig. 1B, GSE271120- Table S1). Furthermore, to determine the early and late <u>d</u>ifferentially <u>e</u>xpressed <u>g</u>enes (DEGs) in response to TRAIL we compared the gene expression at 3h and 24h relative to 0h, by volcano plot (threshold 1 for Log2 fold change, adj p value < 0.001, Fig. 1B, GSE271120-Table S1). The top 25 differentially up-regulated genes at 3h and 24h included the cytokines, CXCL1, CXCL8, CSF2, IL6, CXCL3, IL11, CXCL2, CXCL11 (Table S2). In addition to these cytokines, BIRC3 (that codes for Inhibitor of <u>A</u>poptotic <u>P</u>rotein) and TRAF1 genes were also upregulated. From these DEGs, pathway enrichment analysis was performed using List 2 Pathways (L2P, https://github.com/ccbr/l2p). Analysis platform (NIDAP) which indicated that the Tumor Necrosis Factor-alpha (TNFA) signaling via NFKβ pathway was upregulated at all time points (Fig. 1C). Gene Set Enrichment Analysis (GSEA) performed at all time points also confirmed the hallmark of TNFA Signaling via NFKβ with the highest enrichment score at 3h, when the leading-edge genes were also identified (Fig. 1D). The 10 most up/downregulated pathways identified by GSEA after 3h of TRAIL treatment are shown in Supplementary Fig. 1C.

**Fig. 1.**
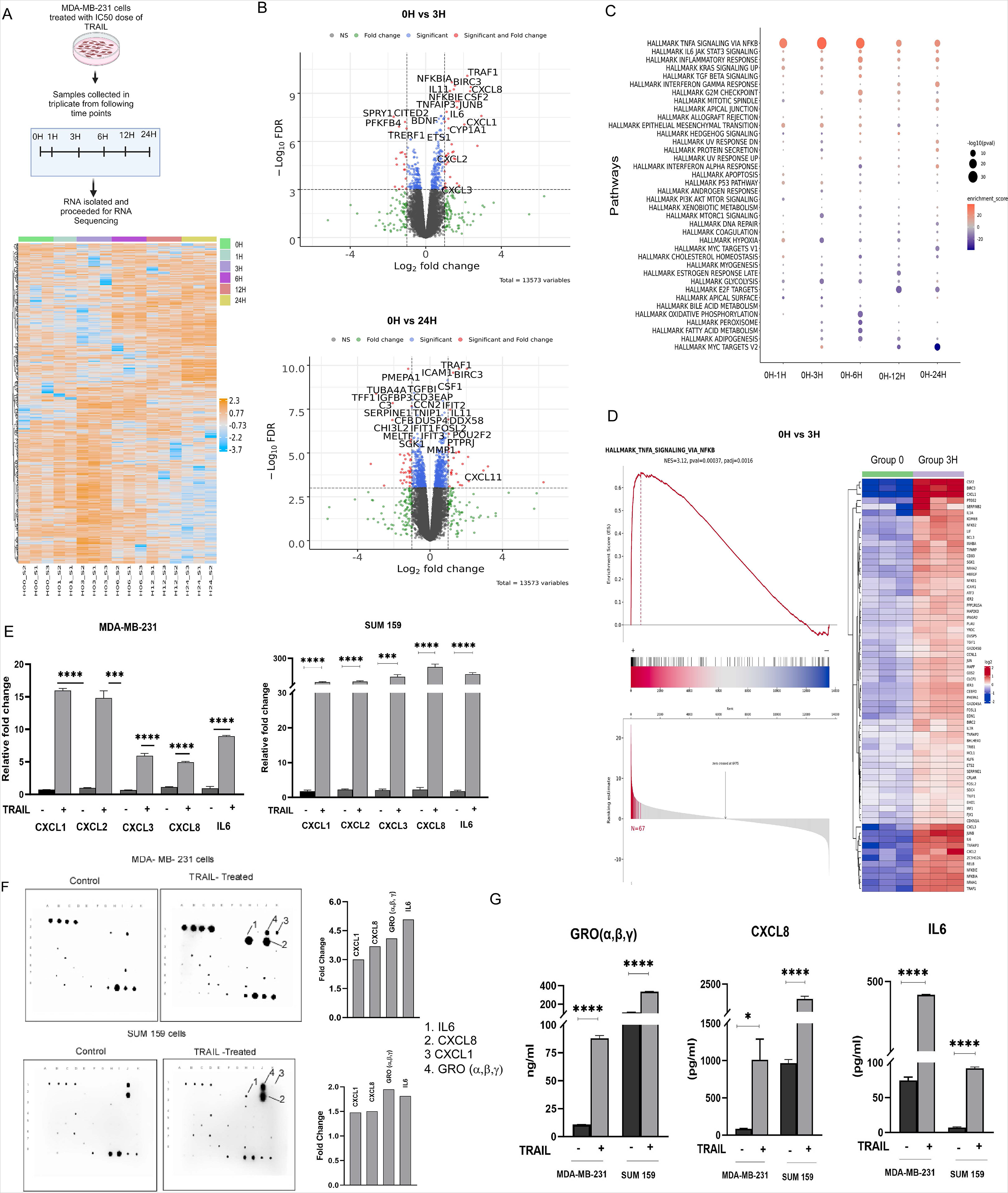
TRAIL upregulates the expression of cytokines in human TNBC cells. A. (Upper panel) Transcriptomic experimental design of TNBC cell line, MDA-MB-231 incubated with GST- TRAIL (45nM). (Lower panel). Heatmap showing upregulated (orange colored) and downregulated (blue colored) genes in MDA-MB-231 cells treated with GST-TRAIL at 0h, 1h,3h, 6h, 12h and 24h in triplicate. Top 500 genes based on variance were used from normalized counts of the RNA- seq data to generate the heat map. B. Upper panel- Volcano plots showing GST- TRAIL-induced upregulated genes at 3h compared with untreated cells (0h), Lower panel-Volcano plots showing GST-TRAIL upregulated genes at 24h compared with untreated cells (0h). Vertical dotted lines indicate Log 2-fold change and horizontal dotted line indicates p value = 0.001. C. Gene set enrichment analysis (GSEA) for the hallmark gene set. D. GSEA enrichment plot and heatmap showing GST-TRAIL-induced gene expression alteration in TNFA signaling via NF-κB. Hallmark pathway at 3h. E. qRT-PCR of MDA-MB-231 and SUM159 mRNA for TRAIL-induced upregulated cytokine genes at 3hdentified by RNAseq. Data are mean +/-SEM of 3 independent experiments. F. Cytokine array with CM or T-CM from MDA-MB-231 (Upper panel) or SUM159 (lower panel) collected after 24h of GST-TRAIL treatment. G. ELISA for indicated cytokines from CM or T-CM of MDA-MB-231 and SUM159 collected after 24h. Data shown as mean +/-SEM of 3 independent experiments. \*\*\*\**p*<0.0001, Student’s *t*-test.

The upregulated cytokines identified by RNA-seq at 3h were confirmed using REAL TIME PCR (qRT-PCR) in MDA-MB-231 and SUM159 cells treated with TRAIL for 3h (Fig. 1E). Consistent with the RNAseq data, qRT-PCR also confirmed that BIRC3 and TRAF1 mRNA levels were significantly increased by TRAIL treatment at multiple time points (Supplementary Fig. 2A). MEDI3039, a potent TRAIL receptor agonist [14], similarly increased expression of cytokines when compared with untreated controls in MDA-MB-231T cells (Supplementary Fig. 2B). Significant increase in cytokine expression was also observed in ER+, HER2+ and additional TNBC breast cancer cell lines, indicating that TRAIL induces cytokine production in all subtypes of breast cancer cells irrespective of TRAIL sensitivity (Supplementary Fig. 2C; Table S3) [13, 14]. Protein expression measured by cytokine array analysis indicated altered expression of various cytokines (Supplementary Fig.3A, B), including CXCL1, GRO (CXCL1/α, CXCL2/β and CXCL3/γ), CXCL8 and IL6 proteins which increased commonly by TRAIL treatment in both MDA-MB-231 and SUM159 cells (Fig. 1F). We further confirmed the TRAIL-induced increase in GRO, CXCL8, and IL6 proteins by ELISA (Fig. 1G).

To examine if TRAIL induces immune modulatory cytokines in TNBC tumors *in vivo,* we harvested orthotopic MDA-MB-231T xenografts, 24h post treatment with or without a dose of the TRAIL agonist Apo2L (Fig. 2A, upper panel). Single dosing of Apo2L reduced the tumor weight at the time of necropsy, although not significantly (Fig. 2A, lower panel). Activation of the TRAIL apoptotic pathway by Apo2L in the tumor tissue was demonstrated by cleaved caspase-8 in treated mice compared to control (Fig. 2B). Apo2L administration induced human cytokine transcripts (Fig. 2C) and human CXCL8 and IL6 protein expression (Fig. 2D) in the tumor lysates along with a significant increase in serum IL-6 levels, but not serum CXCL8 (Fig. 2E).

**Fig. 2.**
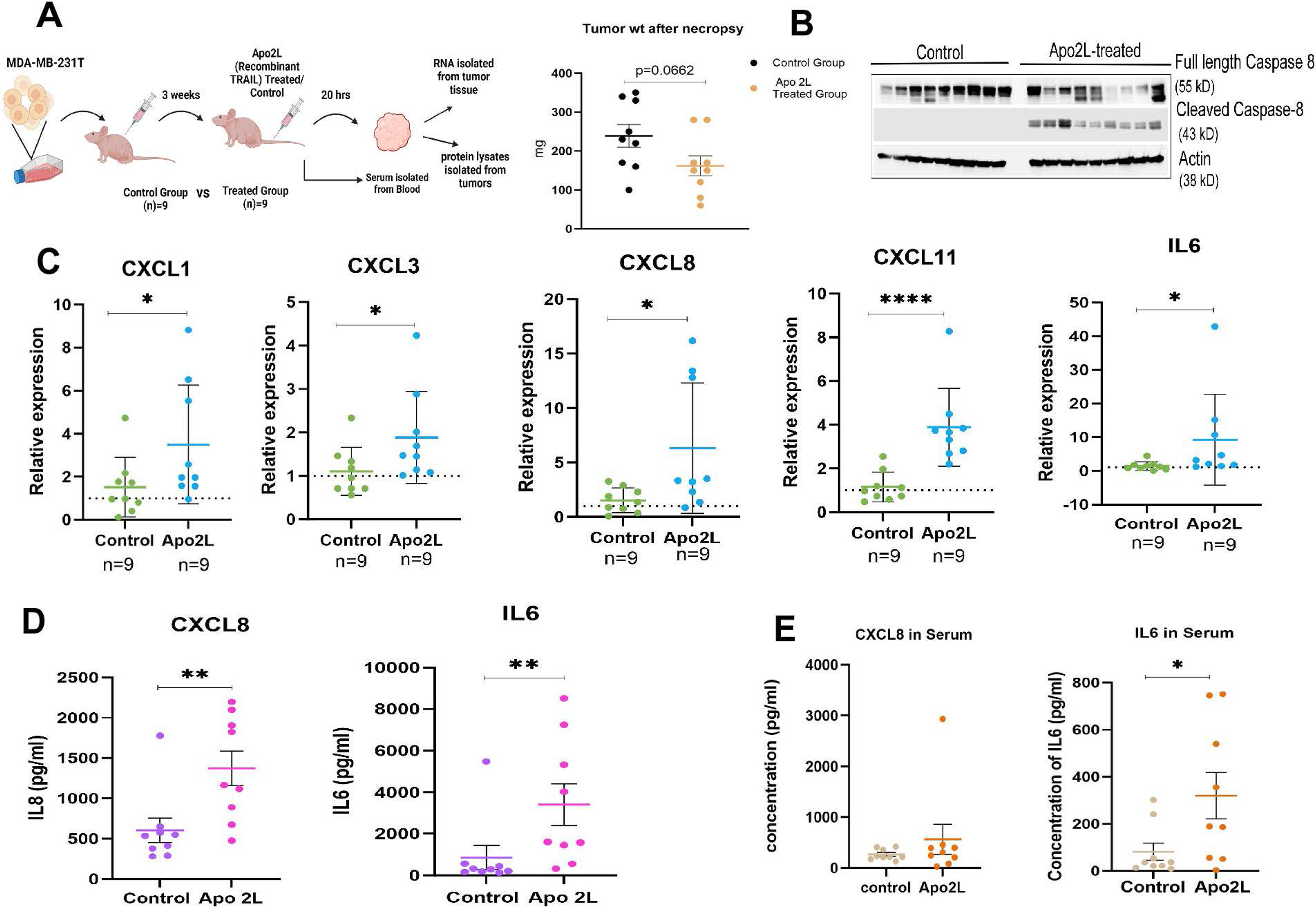
TRAIL induces the cytokine expression *in vivo*. A. Upper Panel-Experimental procedure outline. 5X10^6^ MDA-MB-231T cells were implanted in the mammary fat pad of nude mice. When the tumors reached (200 mm^3^), mice were randomized into two groups of 9 mice each and injected via tail vein with Apo2L (recombinant TRAIL; 100 μg/200ul/ body weight) or control vehicle (PBS). Adjacent panel shows tumor weight in Apo2L treated and control groups 24 h post treatment. B. Activation of TRAIL signaling apoptotic pathway was assessed by immunoblotting for full length and cleaved caspase-8 protein for tumors treated with APO2L or control. C. mRNA expression of human cytokines induced by APO2L vs control from tumors measured by qRT-PCR. D. Protein levels of human CXCL8 and IL6 induced by APO2L vs control from tumor lysates measured by ELISA. E. Serum levels human CXCL8 and IL6 in tumor bearing mice induced by APO2L vs control measured by ELISA. Data are shown as mean +/-SD. \**p*<0.05, \*\**p*<0.01, \*\*\*\**p*<0.0001, Student’s *t*-test.

Taken together, these results confirmed that exogenous addition of TRAIL to human TNBC cells and tumors induces immune modulatory cytokine production *in vitro* and *in vivo*.

### DR5 receptor mediates TRAIL-induced cytokine production via a caspase-activity independent pathway

TRAIL induces apoptosis via ligand binding to the death receptors DR4 and DR5, and activation of caspase-8 [33]. Silencing DR4 and DR5 simultaneously using siRNA, inhibited TRAIL-induced cytokine production (Supplementary Fig. 4A). However, silencing of DR5 compared to silencing of DR4 had a larger effect on TRAIL-induced cytokine production in MDA-MB-231 and SUM159 cells (Fig. 3A and B). Knockdown of DR4 and DR5 was confirmed by both immunoblot and qRT-PCR (Fig. 3C).

**Fig. 3.**
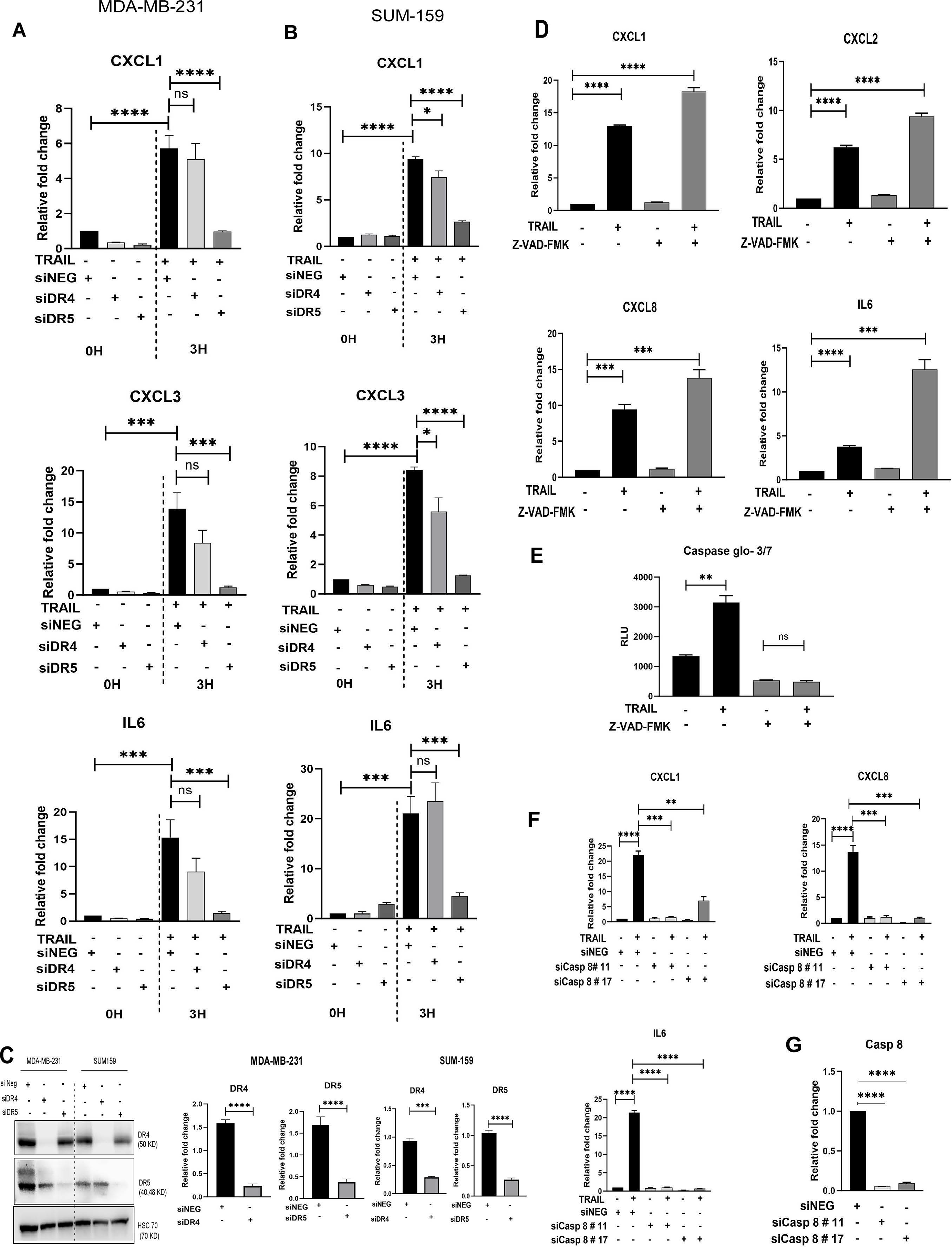
TRAIL induces cytokine mRNA predominantly through DR5 and caspase 8 independent of its caspase activity. A-B. Cytokine mRNA expression in MDA-MB-231 (A) or SUM159 (B) TNBC cells with either siRNA knockdown of DR4 or DR5 after 3h treatment with or without GST-TRAIL measured by qRT-PCR. C- Left panel, immunoblot and right panels- qRT-PCR confirming the knockdown of DR4 and DR5 in indicated cell lines. Data are mean+/- SEM for 3 independent experiments. D. Effect of administration of Z-VAD-FMK on GST-TRAIL-induced cytokine mRNA production in MDA-MB-231 cells measured by qRT-PCR. E. GST- TRAIL induced caspase activity measured by Caspase-Glo 2.0 assay in cells treated +/- Z-VAD-FMK. Data are mean+/- SEM of 3 independent experiments. F. Effect of siRNA knockdown of Caspase 8 on GST-TRAIL-induced indicated cytokines in MDA-MB-231 cells. G. confirmation of the Caspase 8 mRNA knockdown by qRT-PCR. \**p*<0.05, \*\**p*<0.01, \*\*\**p*<0.001, \*\*\*\**p*<0.0001, Student’s *t*-test.

Previously published data using cervical and colorectal cancer cell lines demonstrated that caspase-8 protein but not caspase activity was required for TRAIL induction of cytokines [34]. Therefore, we tested whether inhibition or silencing of caspases suppresses the TRAIL induced cytokine production in TNBC. Pre-treatment with Z-VAD-FMK, a pan-caspase inhibitor, failed to inhibit TRAIL-induced cytokine mRNA increase in TNBC cells despite completely inhibiting TRAIL-induced caspase activation (Fig. 3D, E). Z-VAD-fmk treatment further increased cytokine mRNA induction by TRAIL (Fig.3D, E), likely due to increased viable cells. In contrast, caspase- 8 silencing by siRNA inhibited the TRAIL-induction of cytokine transcripts (Fig. 3F). Knockdown of caspase-8 was confirmed by measuring mRNA levels using qRT-PCR (Fig. 3G).

These findings demonstrate that DR5 and caspase-8 are required for TRAIL-induced cytokine production in a caspase- activity independent manner.

### TRAIL-induced cytokine production in TNBC is mediated by non-canonical NFKB2 pathway

As RNAseq data showed upregulation of multiple components of the NFKB1 and NFKB2 pathway and pathway analysis identified the TNFA/NFKβ-signaling pathway as the most upregulated pathway by TRAIL (Fig. 1C, Supplementary Fig. 1C) we next delineated the role of the NFKB pathway in the TRAIL-mediated cytokine induction. TRAIL treatment of MDA-MB-231 cells induced expression of NFKB1 protein (p105/p50) and phosphorylation of p65 (Fig.4A). Concomitantly, it also increased the phosphorylation of p100 (s866/s870) and expression of NFKB2 protein (p100/p52) in a dose dependent manner (Fig. 4B) suggesting the activation of both canonical (NFKB1) and non-canonical (NFKB2) pathways. Similar increased phosphorylation of p100 (s866/s870) was also observed in other TNBC cell lines (Fig. 4C).

**Fig. 4.**
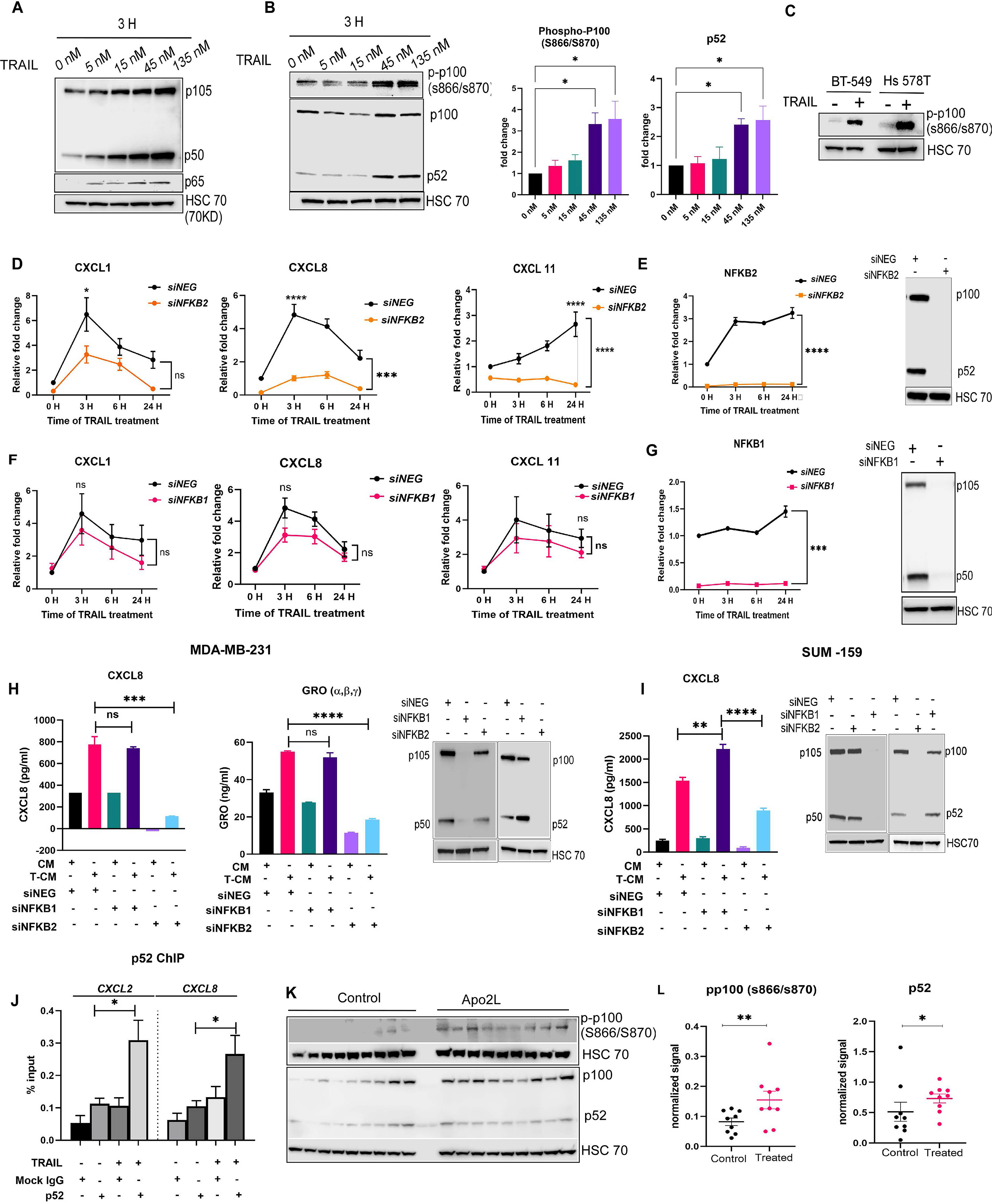
NFKB2 regulates TRAIL induced cytokine production in TNBC. A. Representative immunoblot of NFKB1 (p105/p50) and p65/RELA in MDA-MB-231 cells treated with indicated concentration GST-TRAIL for 3h. B. Left panel-representative immunoblot of phospho-NFKB2 (p-p100) ad expression of NFKB2 (p100/p52) in MDA-MB-231 cells treated with indicated concentration GST-TRAIL for 3h HSC 70 was used as a loading control for both blots. Right panel- quantification of p-p100 and p52 in cells treated with TRAIL at indicated concentrations normalized to HSC 70. Data in graphs are the mean+/- SEM of 3 independent experiments: Student’s t- test. C. Immunoblot of phospho-NFKB2 (p-p100) in in BT549 and Hs 578T cells after 3h after incubation +/- GST-TRAIL. D. Cytokine mRNA levels measured by qRT-PCR in MDA-MB-231 cells with siRNA knockdown of NFKB2 or siNEG control treated with GST- TRAIL for indicated time points, E. Left panel- NFKB2 mRNA level measured by qRT-PCR from cells used for experiments shown in D; Right panel- representative immunoblot of NFKB2 levels from experiments shown in D. F. Cytokine mRNA levels measured by qRT-PCR in MDA- MB-231 cells with siRNA knockdown of NFKB1 or siNEG control treated with GST-TRAIL for indicated time points, G. Left panel- NFKB1 mRNA level measured by qRT-PCR from cells used for experiments shown in F; Data are the mean+/- SEM of 3 independent experiments. 2-way ANOVA was used for the time course comparisons. H. Left panel- protein levels for indicated cytokines in supernatants from MDA-MB-231 cells with siRNA mediated NFKB1 or NFKB2 gene knock down 24h post treatment +/- GST-TRAIL measured by ELISA. Right panel- Immunoblot confirming the knockdown of NFKB1 and NFKB2 in MDA-MB-231 cells. I. Left panel- protein levels for indicated cytokines in supernatants from SUM-159 cells with siRNA mediated NFKB1 or NFKB2 gene knock down 24h post treatment +/- GST-TRAIL measured by ELISA. Right panel- Immunoblot confirming the knockdown of NFKB1 and NFKB2 in SUM-159 cells J. ChIP-PCR assay for p52 binding at the CXCL2 or CXCL8 promotors using non targeting IgG and p52 antibodys in MDA-MB-231 cells treated +/- GST-TRAIL for 2.5h. Data are the mean +/-SEM of 3 independent experiments. K-L. Immunoblot and quantification from tumor protein lysates prepared from tumors treated with APO2L or control (saline) for phospho NFKB2 (p-p100; S866/S870) and p100/p52 protein levels Quantification is shown as the mean +/-SD (n=9 in each group); Student’s *t*-test. \**p*<0.05, \*\**p*<0.01, \*\*\**p*<0.001, \*\*\*\**p*<0.0001.

Next, we investigated the potential role of NFKB1 and NFKB2 pathway activation in the TRAIL-induced cytokine production. Knockdown of NFKB2 but not of NFKB1 in MDA-MB-231 significantly inhibited the increase in mRNA expression of TRAIL-induced cytokine (Fig. 4D-G and Supplementary Fig. 5A, B). Additionally, NFKB2 knockdown but not of NFKB1 significantly inhibited the TRAIL-induced increase of cytokine proteins in cell supernatants from MDA-MB- 231 and SUM159 TNBC cells (Fig. 4H and I, respectively). ChIP assay demonstrated TRAIL-induced binding of p52 to the cytokine promoters for CXCL2 and CXCL8 confirming the activation of nuclear p52 subunit upon TRAIL administration (Fig. 4J). Furthermore, Apo2L- administered xenografts showed a significant increase in phosphorylation of p100 and p52 expression as compared to controls (Fig. 4K, L), corroborating the activation of the NFKB2 pathway induced by TRAIL *in vivo*.

Collectively these data demonstrated that the non-canonical NFKB2 pathway is activated by TRAIL both *in vitro* and *in vivo* and that TRAIL-induced cytokine production primarily depends on NFKB2 expression.

### TRAIL-induced cytokines significantly increase neutrophil chemotaxis

TRAIL-induced CXCL1, CXCL2, CXCL3, CXCL8, IL6 are known to promote neutrophil chemotaxis and activation [35, 36]. Therefore, we examined the chemotactic property of supernatants from TRAIL treated TNBC cells for human neutrophils. Neutrophils isolated and characterized from normal human donor blood (Supplementary Fig. 6A) were incubated in either CM or T-CM (collected as described in the methods from untreated TNBC cells-CM and TRAIL-treated TNBC cells-T-CM). Confocal microscopy showed marked elevation of polymerized actin and increased pseudopodia formation in neutrophils incubated in T-CM compared to those incubated in CM (Fig. 5A). Neutrophil chemotaxis was significantly increased by T-CM compared to CM from several breast cancer cell lines (MDA-MB-231, SUM-159, MDA-MB-453, MCF-7) although the increase was modest when using T-CM from MCF-7 (Fig. 5B, C). Direct addition of exogenous TRAIL to serum free media (SFM) or CM had a minimal effect on neutrophil chemotaxis compared to that of T-CM, indicating that TRAIL-induced factors play a role in neutrophil chemotaxis rather than TRAIL itself (Fig. 5B, right panel). Furthermore, neutrophil chemotaxis was significantly decreased when using T-CM from MDA-MB-231 with NFKB2 knockdown but not from cells with NFKB1 knockdown (Fig. 5D, E), consistent with the NFKB2 dependence for production of the cytokines as shown above in Fig. 4D-I.

**Fig. 5.**
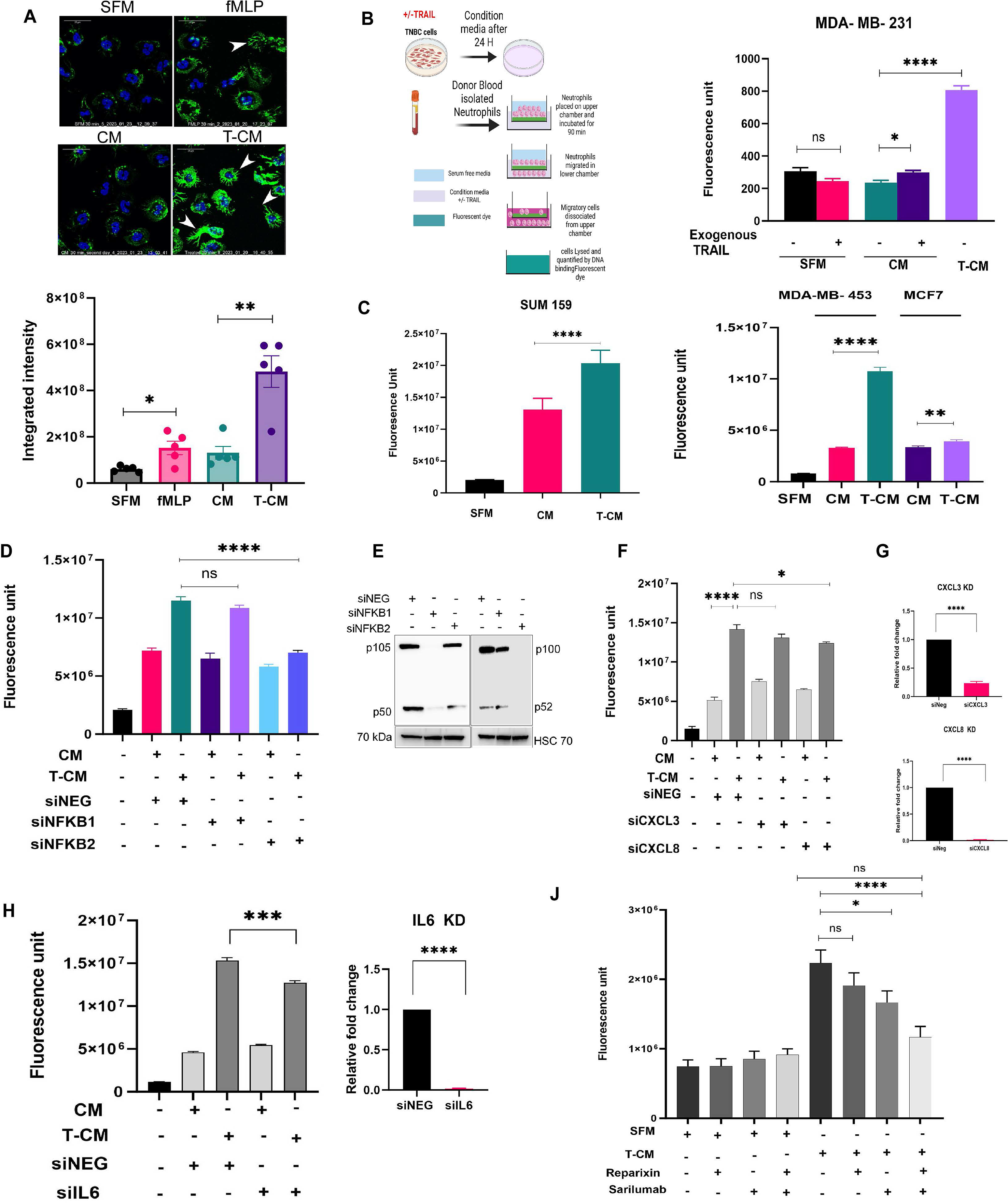
Cytokines produced by TNBC cells in response to TRAIL promote neutrophil chemotaxis in an NFKB2 dependent manner *in vitro*. A. Upper panel-Representative field of immunofluorescence staining of F-actin (green) and DAPI (blue) in neutrophils incubated in either fMLP (positive control), SFM (with DMSO added), CM or T-CM for 30 min. Pseudopodia are indicated as closed arrow heads on image) in neutrophils. Scale bar, 25 μm. Lower panel- Graphical quantification of F-actin intensity within neutrophils incubated in designated condition media. At least 5 random fields were selected for each sample and results are mean +/- SEM from n=5 individual experiments. B. Left panel- schematic representation of the chemotaxis assay with donor blood isolated neutrophils against CM from +/- GST-TRAIL-treated TNBC cells. Right panel- chemotaxis assay of neutrophils to SFM, or CM or T-CM from MDA-MB-231 cells collected 24h post incubation +/- GST-TRAIL. For SFM and CM, TRAIL was added as indicated to examine the effects of TRAIL in the T-CM. C. Left panel- Chemotaxis of neutrophils against indicated SFM, CM or T-CM from SUM159 produced as outlined in B. Right panel- Chemotaxis of neutrophils against indicated CM or T-CM from MDA-MB-453 and MCF7. D. Chemotaxis of neutrophils against CM or T-CM collected from MDA-MB-231 pretreated with siNEG (control), siNFKB1, siNFKB2 gene knockdown. E. Immunoblot confirming the knockdown of NFKB1 and NFKB2 in lysates from MDA-MB-231 cells included in the chemotaxis experiment as shown in D. F. Chemotaxis of neutrophils against CM or T-CM collected from MDA-MB-231 pretreated with siNEG (control), siCXCL3, siCXCL8 gene knockdown. G. Confirmation of confirming the knockdown of indicated mRNA gene by qRT-PCR in cells used in the experiment as shown in F. H. Chemotaxis of neutrophils against CM or T-CM collected from MDA-MB-231 cells pretreated with siNEG(control) or siIL6 gene knockdown. I. Confirmation of IL-6 mRNA knockdown from cells used in H by qRT-PCR. J. Neutrophil chemotaxis to SFM orT-CM from MDA-MB-231 with reparixin (1 μM; CXCR1/2 inhibitor) or sarilumub (200 μM; IL6R inhibitor) treated neutrophils. Data from the chemotaxis assays are mean+/-SEM of 3 independent experiments. \**p*<0.05, \*\**p*<0.01, \*\*\**p*<0.001, \*\*\*\**p*<0.0001, Student’s *t*-test.

CXCL (1, 2, 3, 8) chemokines mediate their signal either via C-X-C chemokine receptor type 1 (CXCR1) and/or C-X-C chemokine receptor type 2 (CXCR2) on the neutrophils [37, 38].

Silencing CXCL (1, 2, 3, 8) either individually, or in combination in the TRAIL treated TNBC cells or blocking CXCR1/R2 on neutrophils with Reparixin, an allosteric inhibitor [39] partially inhibited the T-CM mediated neutrophil chemotaxis (Fig. 5F, G, J, Supplementary Fig. 6B-D). Similarly silencing IL-6 in the TRAIL treated TNBC cells or inhibiting IL-6 receptors on the neutrophils with Sarilumab, an IL-6 receptor antagonist [40] only partially reduced the T-CM mediated neutrophil chemotaxis (Fig. 5H-J, Supplementary Fig. 6D). Notably, incubating the neutrophils with both Reparixin and Sarilumab reduced T-CM induced neutrophil chemotaxis more than either drug alone thus implying that the T-CM induced increased chemotaxis of neutrophils is meditated by multiple TRAIL-induced cytokines (Fig. 5J).

### TRAIL induces neutrophil recruitment *in vivo*

We used several orthotopic TNBC xenograft models to investigate TRAIL-induced recruitment of neutrophils to tumors in vivo. Tissue lysates from the orthotopic xenografts described above (Fig. 2A) demonstrated a significant TRAIL-induced increase of Ly-6G expression in the tumor tissue consistent with neutrophil accumulation (Fig. 6A). Neutrophil infiltration (CD45^+^ Ly-6G^+^ cells) in orthotopic xenograft TNBC tumors was also increased by the TRAIL agonist MEDI3039 [14] when compared to control tumors visualized using CODEX analysis (Fig. 6B, C, Supplementary Fig. 7). Additionally, IHC indicated that myeloperoxidase (MPO), a lysosomal protein abundantly expressed in neutrophils was significantly increased in MEDI3039 treated tissues compared to control (Fig. 6D), further supporting that TRAIL promotes neutrophil infiltration into tumors. Intravital imaging of dendra-2-labelled MDA-MB-231 tumors 24h post Apo2L injection demonstrated increased recruitment of labelled neutrophils when compared to saline treated control tumors. (Fig. 6E, F). These data demonstrated that TRAIL treatment promotes neutrophil recruitment to tumors *in vivo*.

**Fig. 6.**
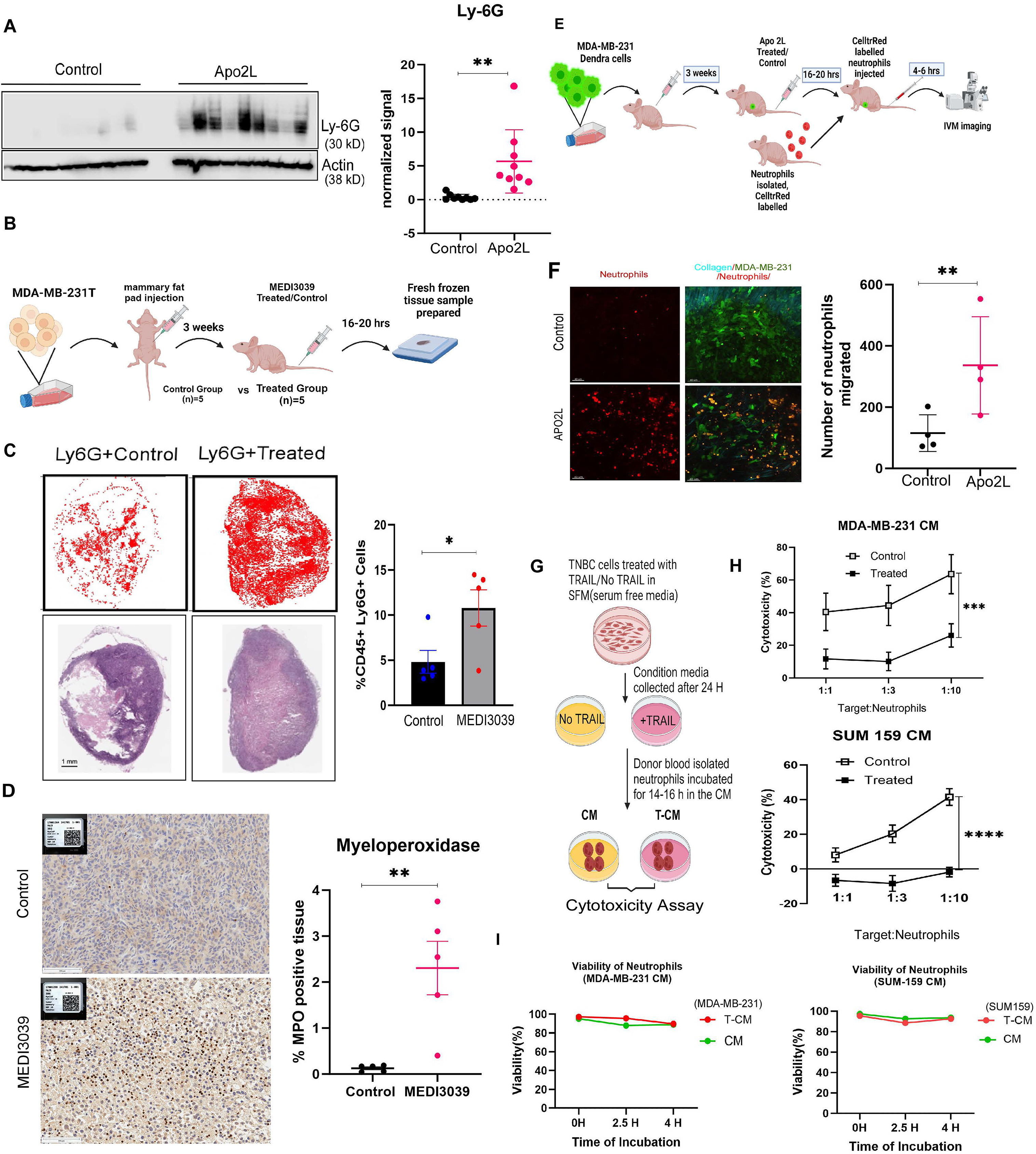
APO2L injection recruits neutrophil *in vivo* and modifies the function of neutrophils. A. Lysates from tumors (described in Fig. 2A) treated +/- APO2L (100 μg/200 μl /body weight) were immunoblotted for the murine neutrophil marker LY-6G (Left Panel) and quantified for expression of Ly-6G (right panel). B. Experimental design for assessing neutrophil recruitment in response to TRAIL agonist (MEDI3039, 0.3 mg/Kg) in tumor tissue by CODEX. C. Upper panel- Spatial dot plot of CODEX image and analysis from a representative sample showing recruitment of LY-6G+ cells in tumor tissue from mice injected +/- MEDI3039. Lower panel- hematoxylin and eosin staining of the same tissue. Right panel- Quantification of the % of CD45+LY-6G+ cells in the tumors treated +/- APO2L (n=5 in each group). D. Myeloperoxidase (MPO) immunohistochemistry analysis of tumor samples from MEDI3039or vehicle-injected mice. Scale 100 μm. E. Experimental design for assessing neutrophil recruitment in response to APO2L (100μg /200μl/body weight) compared to vehicle- injected mice (n=4) by intravital microscopy imaging. F. Representative intravital image of cell tracker red stained neutrophils in tumors in mice treated with saline vehicle (upper panels) or+/- APO2L (lower panels). Neutrophils alone are shown in red (left panels) and merged images (right panels) showing neutrophils (orange), dendra- 2 labelled MDA-MB-231 tumor (green) and collagen (cyan) are shown in the right panels. G. Quantification of number of neutrophils recruited by for tumors treated +/- APO2L (N=4 for each group) Data in A, C, D and F are mean+/-SD. \**p*<0.05, \*\**p*<0.01 Student’s *t*-test. G. Schematic representation demonstrating the conditions for *in vitro* cytotoxicity assay of neutrophils. H. Killing of Luciferase (Luc) labeled LM2- MDA-MB-231cells cocultured with neutrophils incubated in either CM or T-CM from MDA-MB-231 cell or SUM 159 as indicated. I. Viability of neutrophils analyzed with AOPI assay over the time frame of the cytotoxicity experiment. Data are mean +/-SEM using 2-way ANOVA for n=5 independent experiments, \*\*\**p*<0.001, \*\*\*\**p*<0.0001.

### T-CM suppress neutrophil cytotoxicity against tumor cells

Neutrophil-mediated anti-tumor cell cytotoxicity [41–43] was suppressed by incubating neutrophils in T-CM compared to CM from two different TNBC cell lines (Fig. 6G, H). AOPI staining demonstrated no significant loss of viability of neutrophils incubated in T-CM or CM over the time of the assay (Fig. 6I), indicating that the impaired neutrophil cytotoxicity by T-CM was not due to the reduced cell viability of neutrophils. These findings show that T-CM of TNBC cells has a suppressive effect on effector capacities of human neutrophils.

### T-CM and TRAIL alter the neutrophil transcriptome

To further characterize the effect of T-CM on neutrophils, we performed bulk RNAseq of healthy donor-derived, isolated neutrophils incubated in either SFM, SFM-T (SFM with addition of exogenous TRAIL), CM or T-CM. Principal component analysis and unsupervised clustering highlighted a clear segregation of transcriptomes between the four conditions. Notably, there was a profound reprogramming of the transcriptome when comparing SFM-T to SFM, establishing a direct effect of TRAIL on the neutrophil transcriptome (Supplementary Fig. 8A, Fig. 7A). Differential gene analysis indicated that 896 genes (T-CM vs CM) and 111 genes (SFM-T vs SFM) were altered within the designated groups. Venn diagram showed an overlap of 91 genes among SFM-T vs SFM, T-CM vs CM and T-CM vs SFM (Fig. 7B). Analysis of DEGs revealed significant upregulation of several immunoregulatory cytokines and inflammatory genes (e.g., CXCLs 2, 3, CCLs 1, 4, &, interleukins 1α, 1β, & 6) and MMPs (9,14) in both T-CM and SFM-T incubated neutrophils, when compared to CM and SFM respectively (Fig. 7C, GSE271122-Table S4).

**Fig. 7.**
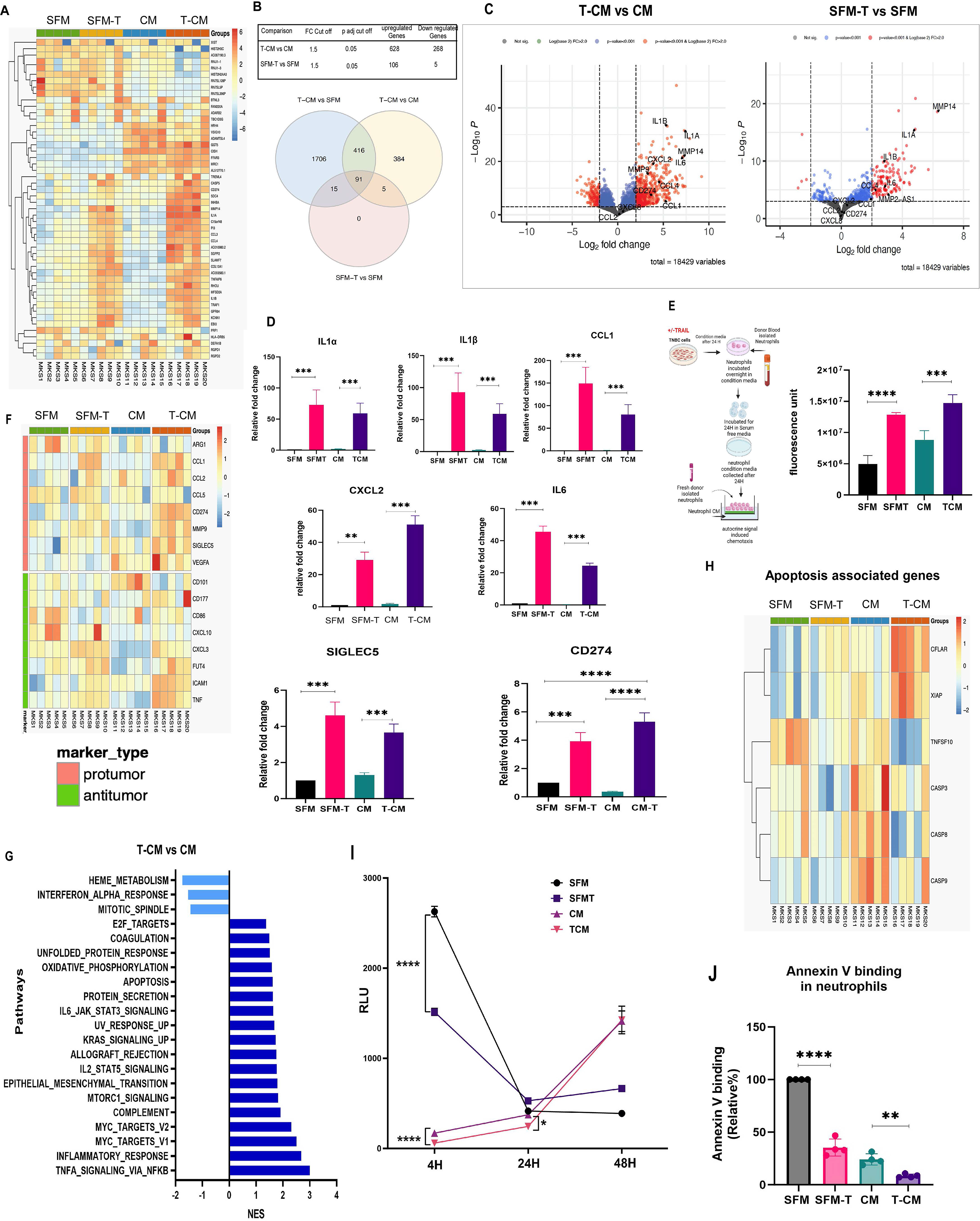
T-CM and TRAIL modify the transcriptomic profile of neutrophils. A. Heat map describing 50 genes with highest variance across neutrophil samples isolated from n=5 donors and incubated in SFM, SFM-T, CM, and T-CM for 18h. B. Table showing number of genes upregulated and down regulated in T-CM vs CM and SFM-T vs SFM. Venn diagram shows <u>d</u>ifferentially <u>e</u>xpressed gene overlap between the indicated sample groups at fold change threshold 1.5, padj cutoff:0.05. C. Volcano plots showing differential analysis for T-CM vs CM and SFM-T vs SFM. D. qRT-PCR validation of representative DEG’s that are significantly upregulated in indicated groups. Data shown are mean+/- SEM of 3 independent experiments. E. Left panel- Experimental design illustrating chemotaxis assay of fresh isolated neutrophils against ‘neutrophil secreted factors.’ Supernatants from neutrophils incubated in SFM, SFMT, CM and T-CM from MDA-MB-231 were prepared as described in methods section. Right panel-quantification of neutrophil chemotaxis to supernatants from the neutrophils treated as indicated. F. Heat map presenting protumor and anti- tumor gene expression in neutrophils treated with SFM, SFMT, CM and T-CM as indicated. G. GSEA for the hallmark gene sets for neutrophils treated with T-CM vs CM. H. Heatmap showing DEG’s associated with apoptosis of neutrophils within indicated group. I. Time- course of Caspase 3/7 activity in neutrophils incubated in different condition media. Data are mean+/- SEM of 3 independent experiments. J. Annexin V binding measured as relative percent compared to SFM incubated neutrophils in neutrophils incubated with indicated conditions for 24h. Data are mean+/- SEM of 3 independent experiments. Comparisons in this figure were analyzed using Student’s *t*-test; \**p*<0.05, \*\**p*<0.01, \*\*\**p*<0.001, \*\*\*\**p*<0.0001.

Alterations in expression of selected genes was confirmed by qRT-PCR (Fig. 7D, supplementary Fig 8B). Additionally, a cytokine array analysis performed with ‘neutrophil condition media’ (collected as described in the methods from CM and T-CM incubated neutrophils) detected increased expression of CXCL8, M-CSF, TGFβ2 and TIMP2 in response to T-CM compared to CM from the SUM-159 cell line (Supplementary Fig. 9A, B). Functionally, conditioned media from neutrophils incubated in SFM-T and T-CM compared to SFM and CM increased neutrophil chemotaxis of untreated neutrophils, suggesting that neutrophils treated with TRAIL and T-CM can in turn recruit additional neutrophils (Fig. 7E).

Exposure of neutrophils to TRAIL and T-CM also induced upregulation of the immune inhibitory checkpoint genes CD274 (PDL1) and SIGLEC-5 [44] (Fig. 7D, Supplementary Fig. 9C). To further evaluate the effects of TRAIL (e.g., SFM-T) and T-CM on the neutrophils we evaluated expression of published protumor and anti-tumor neutrophil marker genes [45–47] (Fig. 7F). TRAIL and T-CM induced both pro- and anti-tumorigenic neutrophil transcriptome changes (Fig. 7F, 7D, Supplementary Fig.8,9). GSEA pathway analysis comparing T-CM vs CM incubated neutrophils and SFM-T vs SFM group indicated TNFA via NFKB pathway to be enriched, analogous to the results in TRAIL treated TNBC cells (Fig. 7G and Supplementary Fig. 9D).

Since TRAIL activates an apoptosis signaling pathway, we analyzed the expression of genes related to apoptosis in neutrophils treated with SFM, SFM-T, CM and T-CM. Expression of the pro-apoptotic genes Casp3, Casp8, and TNSF10 were decreased in T-CM treated neutrophils, while the anti-apoptotic genes CFLAR and XIAP were increased (Fig. 7H). We examined a time course of caspase3/7 activity in neutrophils incubated with SFM, SFM-T, CM and T-CM. At 4h, caspase-3/7 activity was significantly lower in neutrophils incubated in T-CM> CM>>SFM-T compared to those incubated in SFM (Fig 7I). In SFM and SFM-T, caspase activity was highest at 4h and then decreased at 24 and 48h. In contrast, both CM and T-CM had low caspase activity at 4 hours and 24 hours which increased at 48h. Caspase activity was significantly lower in neutrophils incubated in T-CM compared to those incubated in CM at 4 and 24h. (Fig. 7I). Annexin V binding to the neutrophils (a marker of early apoptosis) [48] decreased at 24h by T-CM > CM > SFM-T compared to SFM (Fig. 7J). These data suggest that TRAIL induced delayed-apoptosis in neutrophils.

Overall, exposure of neutrophils to TRAIL or T-CM in tumor cells leads to substantial transcriptomics changes in neutrophils, which may support a pro-tumorigenic phenotype.

### TRAIL-treated neutrophils inhibit CD4^+^T cell proliferation and increase expression of immunosuppressive genes in Tregs

To investigate TRAIL’s effect on adaptive immune cells, we tried to identify an immune competent TNBC mouse model that recapitulated the effects of TRAIL on human TNBC cells. Though some studies report that murine TNBC cells are sensitive to TRAIL [49, 50], we tested 7 murine TNBC cell lines and only the AT3 cell line reached an IC50 at ∼1000 nM (Supplementary Fig. 10A, B); suggesting murine TNBC are relatively resistant to TRAIL compared to the human TNBC cell lines [14]. TRAIL-induced cytokines were observed at the mRNA level in different murine TNBC cell lines but at relatively high TRAIL concentrations (Supplementary Fig 11A). Similarly, a very high dose of TRAIL (1000-5000 nM) caused increased chemotaxis of murine neutrophils against T-CM from MET-1 and AT-3 (Supplementary Fig. 11B). These data suggest that TRAIL does not elicit similar effects on murine TNBC compared to human models. Moreover, TRAIL doses needed to elicit any effects would not be practically achievable in animal models.

Hence, our current study is limited to *in vitro* experiments for exploring the effect of TRAIL-treated neutrophils on other immune cells.

Our finding that TRAIL or T-CM-exposed neutrophils produce different immune modulatory cytokines like CCL (1,2,4), IL (1α, 1β, & 6) (Fig. 7C&D) suggests that these may affect T cell function [51–57]. Thus, we investigated the effect of neutrophils incubated with SFM, SFM-T, CM and T-CM on T cell proliferation. Greater inhibition of T cell proliferation was observed by CM and T-CM incubated neutrophils as compared to SFM or SFM-T incubated neutrophils (Fig. 8A, B). As TRAIL and T-CM significantly upregulated CCL1 chemokine (a ligand for CCR8) expression in neutrophils isolated from healthy human donors (Fig. 7D), we also evaluated the effect of factors secreted from neutrophils incubated with T-CM on the expression of suppressive markers on Tregs (i.e., CCR8) [58–60]. Neutrophil condition media (collected as outlined in the methods from SFM-T and T-CM incubated neutrophils) significantly increased the transcript levels of CCR8 and FOXP3 in isolated Tregs compared to that of SFM and CM (Fig. 8C, Supplementary Fig. 12). The data suggest that neutrophils incubated with T-CM as well as TRAIL potentiate the suppressive phenotype of Tregs. Collectively, our data implies an immune suppressive role of both TRAIL and T-CM incubated neutrophils.

**Fig. 8.**
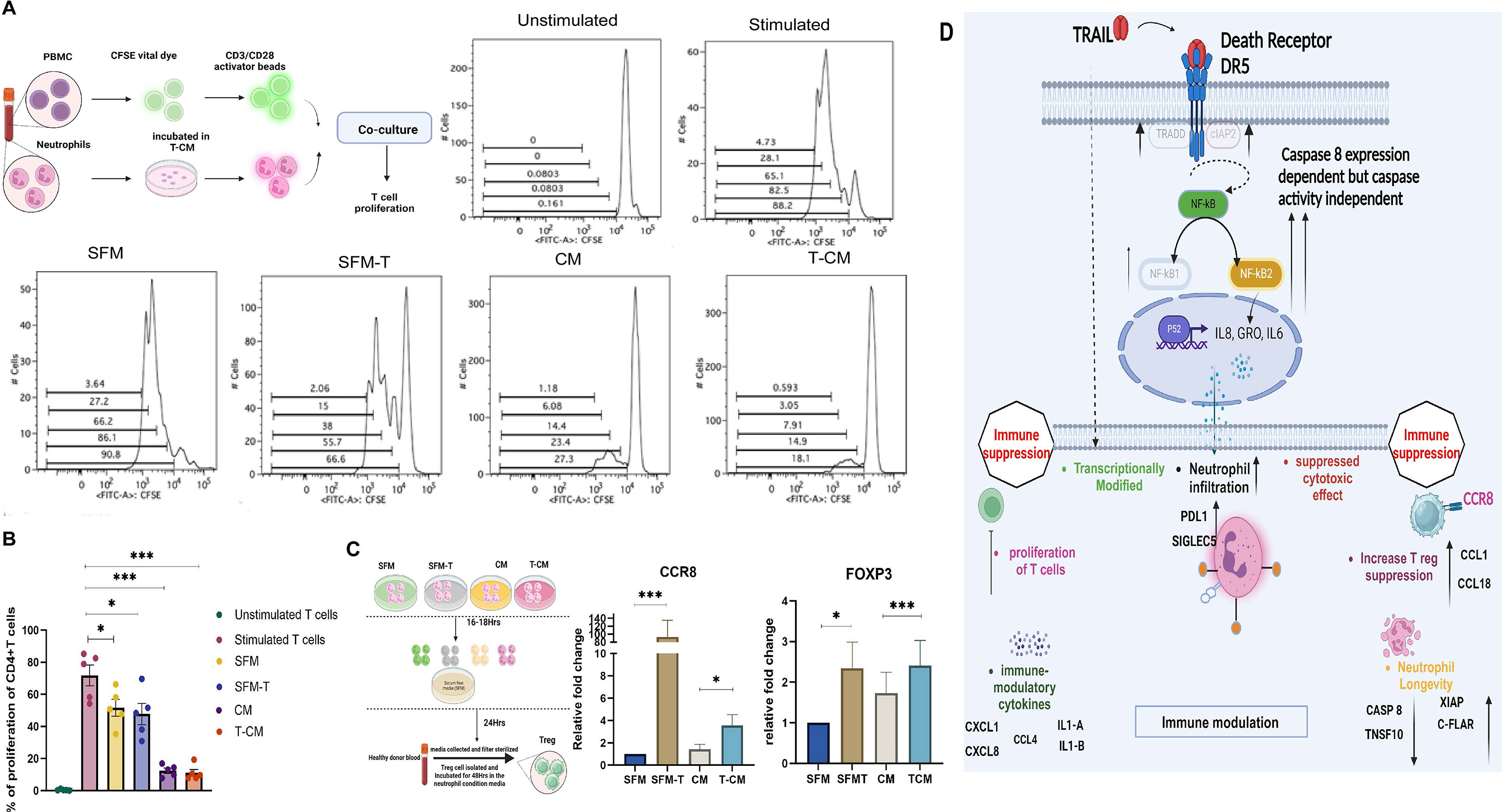
**T-CM and CM incubated neutrophils suppress proliferation of CD4+T cells and promote expression of immune suppressive genes in Tregs**. A. Left- Schematic illustration of proliferation assay of T cells after coculture of PBMC with autologous neutrophils pre-incubated in specified conditions. Representative histograms showing CFSE dilution in CD4+ T cells cultures either unstimulated or stimulated with CD3/CD28 activator beads (right), or stimulated with CD3/CD28 activator beads and treated with neutrophils previously incubated in different conditions (bottom). B. Quantitation of CD4+ lymphocyte proliferation from n=5 experiments. Data shows the percentage of CD4+T cells that proliferated without or with co cultured neutrophils previously incubated in indicated conditions. Data are mean +/- SEM. C. Schematic illustration for donor isolated Treg incubated in designated neutrophil condition media followed by RNA isolation and qRT-PCR. Right panel shows the mRNA expression of CCR8 and FOXP3 in Tregs incubated in specified neutrophil condition media. Data are the mean +/- SEM. D. Graphical model showing that TRAIL induces cytokine expression in TNBC cells via DR5 and Caspase 8in a caspase activity independent pathway via the NFKB2 pathway. These secreted cytokines augment neutrophil recruitment in tumor microenvironment which are transcriptionally modified with immune suppressive function along with altered longevity. Additionally, it highlights TRAIL also alters the neutrophil transcriptome promoting the modification of the immune landscape of tumor microenvironment, ultimately resulting in TRAIL responsive immune modulation. Comparisons in this figure were analyzed using Student’s *t*-test; \**p*<0.05, \*\*\**p*<0.001.

## Discussion

The TRAIL apoptotic pathway is an interesting therapeutic target for the treatment of TNBC exhibiting a selective and robust killing of tumors in many preclinical models [18] . However, TRAIL has not shown significant activity in clinical trials indicating that additional regulatory mechanisms which inhibit TRAIL activity remain elusive [18]. In this study, we evaluated the transcriptional changes in TRAIL-sensitive human TNBC cells induced by TRAIL. We identified multiple genes regulated by TRAIL including multiple immune modulatory cytokines *in vitro* and *in vivo* which were induced predominantly via DR5 and required caspase-8 protein. However, their induction was independent of caspase activity. Our data are consistent with the findings by Martin *et al.*, who reported that TRAIL treatment in cervical and colorectal cancer elicited proinflammatory cytokines in a caspase-independent manner, though caspase-8 was required as a scaffold [34].

GSEA analysis indicated that the TNFα/NFKB pathway was the most enriched pathway by TRAIL treatment of the TNBC cells at all investigated time points. The NFKB family of transcriptional regulators play an important role in modulating tumorigenesis, immune disorders, and response to cancer therapeutics [61]. NFKB signaling can be classified broadly into canonical (NFKB1) and non-canonical pathways (NFKB2). The canonical pathway, mediated by p105/p50 (NFKB1) is typically activated by inflammatory signals which results in the nuclear translocation of p50/RelA dimers where they mediate transcription [62]. The non-canonical NFKB2 pathway activation involves processing of NFKB2 precursor protein p100 to the p52 protein, followed by translocation of p52/RelB dimers to the nucleus where they mediate transcription [63]. Site-specific phosphorylation of serine-866 and serine-870 on p100 is crucial for effective processing of p100 in this alternative pathway. The phosphorylation of p100 at this specific site is dependent on NIK and IKKα [64]. The NFKB pathway has been reported to be involved in TRAIL-induced cytokine production in different cell lines [23]. Our findings confirmed that TRAIL induces the expression of p105/p50 and phosphorylation of p65/RelA (the NFKB1 pathway) in a dose-dependent manner. Furthermore, we report here for the first time that TRAIL induces activation of the non-canonical NFKB2 pathway as measured by increased expression and phosphorylation of p100 (serine-866/870) along with nuclear binding of p52 to the promoter region of target cytokines. Interestingly, silencing of NFKB2 significantly inhibited the production of TRAIL-induced cytokines, whereas silencing of NFKB1 had minimal effect on cytokine expression, suggesting an important role of non-canonical NFKB pathway in the TRAIL therapy.

The upregulated cytokines induced by TRAIL administration in TNBC cells (CXCL1, CXCL2, CXCL3, CXCL8, IL-6) are known regulators of neutrophil function [65, 66]. Neutrophils are the primary responders of the innate immune system and play a dichotomous role in cancer progression as they can mediate tumor cytotoxicity and/or secrete cytokines that can enhance or inhibit anti-tumor immunity [67, 68]. As tumors progress, neutrophils frequently acquire a pro-tumorigenic/immunosuppressive phenotype [27]; moreover, a high neutrophil to lymphocyte ratio is a poor prognostic indicator in TNBC[31]. Our findings demonstrating enhanced neutrophil chemotaxis and recruitment in the TNBC tumor in response to TRAIL suggest a potential role of these neutrophils in modulating the tumor immune milieu. Furthermore, silencing individual cytokines and blocking their receptors by Reparixin [39] and Sarilumab [40] showed only a partial inhibition on chemotaxis of neutrophils in response to T-CM, demonstrating the combined contribution of these cytokines to the neutrophil chemotaxis. Additionally, we found that the neutrophils incubated in T-CM from TNBC cells suppressed cytotoxicity of neutrophils towards the cancer cells implying modification in neutrophil function in response to supernatants from TRAIL-treated-TNBC cells.

The transcriptomic analysis of neutrophils incubated SFM-T as well as T-CM, identified upregulation of multiple cytokines (*e.g.*, CXCL1, CXCL2, CXCL8, IL1α, IL1β, IL-6, CCL1, CCL2, CCL4, TGFβ), proteases (e.g., MMP9, MMP14), and immune-checkpoint proteins (PDL1, SIGLEC5) in the neutrophils which could potentiate tumorigenesis [27, 32, 44]. Furthermore, the induction of cytokine secretion by the neutrophils promoted chemotaxis of additional neutrophils, which is consistent with a paracrine signal relay by the recruited neutrophils [69]. Additional analysis identified altered expression of apoptotic genes (TNSF10, Caspases, CFLAR, XIAP) in TRAIL as well as in T-CM incubated neutrophils, indicating reduced apoptosis. Our result with Caspase 3/7 activity and Annexin V-binding confirmed decreased/delayed caspase activity and apoptosis in neutrophils exposed to TRAIL, CM and T-CM.

*In vitro* studies on T cells and Tregs supported an immunosuppressive role of neutrophils exposed to supernatants of TNBC. Our data shows that neutrophils exposed to CM or T-CM both suppressed T-cell proliferation to a similar degree. Notably, in our *in vitro* experiments we used equal numbers of CM or T-CM treated neutrophils. However, our *in vivo* data demonstrating increased accumulation of neutrophils in tumors in response to TRAIL suggests there would be greater suppression of T cell proliferation due to a greater number of neutrophils. Furthermore, our data indicates an increase in suppressive markers in Tregs in response to secreted factors from neutrophils incubated with TRAIL or T-CM. These findings are in line with previously published work showing that exogenous or endogenous TRAIL induces an immunosuppressive tumor microenvironment [24, 25]. Our study is limited to *in vitro* experiments for exploring the effect of TRAIL-exposed neutrophils on other immune cells as both human and murine TRAIL failed to induce apoptosis or elicit biologically active cytokines in murine TNBC at doses that could be reasonably used in immune competent *in vivo* models.

In conclusion, our findings propose a model whereby TRAIL induced-cytokine production by TNBC via NFKB2 activation recruits neutrophils that potentiate tumor immune suppression (Fig. 8D). These neutrophils are transcriptionally modified, expressing cytokines and immune checkpoint markers in the TRAIL treated-tumor environment, resulting in suppressed cytotoxicity towards cancer cells, and increased the longevity of the neutrophils. Our findings uniquely describe the direct effect of TRAIL on the neutrophil’s transcriptome and production of immunosuppressive cytokines. This implies that TRAIL itself may systematically alter the immune system independent of its effects mediated though the tumor. This study highlights the complexity of crosstalk between TRAIL, TNBC cells, neutrophils, and adaptive immune cells. Our data suggests future therapeutic strategies for TNBC that combine TRAIL agonists with drugs inhibiting the NFKB2 pathway and/or cytokine function.

## Methods

### Reagents

The recombinant protein GST-TRAIL was prepared in the laboratory as previously reported [13]. I*nvitro* experiments were performed with GST-TRAIL (TRAIL) and APO2L (commercially available recombinant TRAIL) was used for in vivo experiments. The details for reagents, equipment and software are listed in Table S5.

### Cell Culture and transfection

The different human and murine breast cancer cell lines and their culture condition used in the current study are described in Table S3. Cell lines authentication was performed using Promega GenePrint 10 system and Mycoplasma tests were routinely conducted in the laboratory using Mycoplasma PCR detection kit.

All RNAi transfection was performed using Lipofectamine RNAiMax following the forward transfection protocol as recommended by manufacturer. Forty-eight hours after transfection, the cells were replated in serum-free media (SFM) followed by GST-TRAIL treatment at the indicated concentration for either 3hours (qRT-PCR), or 24hours (Immunoblotting). To observe the effect of gene knockdowns on cytokine production and chemotaxis, supernatant was collected 24h post TRAIL treatment and used in ELISA and chemotaxis assays. The list of siRNAs used are indicated in Table S5.

### Sample collection and processing from healthy blood donors

Human whole blood samples were obtained from healthy donors on an IRB-approved NIH protocol (99-CC-0168). Neutrophils, and Tregs were isolated from whole blood using the kits and reagents mentioned in Table S5 according to the manufacturer’s protocol. PBMC was isolated by layering of whole blood on density gradient medium using reagents mentioned in Table S5 according to the manufacturer’s protocol. Isolated neutrophils from healthy human donors were first characterized by Flow cytometry with antibodies for CD66B and CD45.The Tregs isolated from fresh PBMC were also characterized by CD45, CD4 and FOXP3 antibodies according to manufacturer’s protocol (Table S5).

### Condition media collection

Multiple human as well as murine TNBC cell lines (Table S3) were either untreated (CM) or treated (T-CM) with indicated dose of human GST-TRAIL (45 nM) or murine- TRAIL for 24h at 37℃ in SFM. The media was then collected followed by centrifugation and filtration (0.22µm) to eliminate debris and stored at -80℃ until use.

In some cases, to address the effect of TRAIL in the T-CM, TRAIL (45nM) was added to the SFM prior to use (SFM-T).

To observe the effect of neutrophil secreted factors on immune cells, isolated neutrophils from healthy donors were incubated in either SFM, SFM-T, CM or T-CM for 16-18h. After the incubation, cells were washed twice with PBS followed by a second round of incubation in SFM for additional 24h. This media containing the neutrophil secreted factors is designated as ‘neutrophil condition media’ which were then collected followed by centrifugation and filtration (0.22µm) to eliminate debris and stored at -80℃ until use.

### Acridine orange/Propidium iodide (AO/PI) cell viability assay

Live/dead, total cell numbers were counted with AOPI and the Cellometer K2 (Table S5).

### RNA extraction and quantitative Real-time PCR (qPCR)

Total RNA was extracted from cell lines, immune cells from donors, or tissues using RNeasy Mini Kit or Trizol followed by qRT-PCR per the manufacturer’s protocol. Isolated Tregs and T effectors were incubated in ‘neutrophil condition media’ for 48hours followed by RNA isolation. Relative quantification of gene expression was normalized to GAPDH or Actin (using the 2^−ΔΔCt^ method). The list of primers used are provided in Table S5.

### RNA- Seq and bioinformatic data analysis

MDA-MB 231 cells were treated with TRAIL (45 nM) for different time points in triplicate followed by RNA isolation, quality control check, and cDNA library preparation. Downstream analysis and visualization were performed using the NIH Integrated Analysis Platform (NIDAP) using R programs developed by the team of NCI bioinformaticians on the Foundry platform (Palantir Technologies). Briefly, RNA-Seq FASTQ files reads were aligned to the reference genome (hg38) using STAR [70] and raw counts data produced using RSEM [71]. The gene counts matrix was imported into the NIDAP platform, where genes were filtered for low counts (<1 cpm) and normalized by quantile normalization using the limma package [72]. Differentially expressed genes were calculated using limma-Voom [73]. Heatmaps was generated using normalized RNA seq data based on the variance. Volcano plots of different time points were generated from DEG analysis compared to 0h (log2fold change ≥1; significant p value of 0.001). The code is available at the following location: https://github.com/NIDAP-Community/TRAIL-Induced-NFKB2-Pathway-Promotes-Neutrophil-Chemotaxis-and-Immune-Suppression-in-TNBC.

Differential gene expression analysis was also performed with (n= 5) individual healthy donor isolated neutrophils incubated for 16 to 18h in either SFM, SFM-T, CM or T-CM. DEseq2 (v1.42.0) [74] software with the RSEM counts data was used for analysis of neutrophil RNA seq. Read counts were filtered to retain genes with at least 10 counts in the smallest group. Gene set enrichment analysis (GSEA) for the RNA-seq data was performed with GSEA 4.3.2 [75] and MSigDB [76].

RNA-Seq data are available through Gene Expression Omnibus (GEO) (accession numbers GSE271120 for MDA-MB-231 RNA seq and GSE271122 for neutrophil RNA seq).

### Cytokine array and ELISA

To validate both the RNAseq data (TNBC cell line and neutrophils) at protein level expression, the Human Cytokine Antibody Array C5 (Table S5) was used. Semi-quantitative detection of 80 proteins in culture supernatants were used according to the manufacturer’s instructions. Signal intensity on the membranes were visualized on an Odyssey, using image studio ^TM^ software. The relative quantity of each protein present in either untreated or treated samples was normalized to the positive controls included on the array. Results were expressed as fold change of each factor in the treated sample compared toto the untreated sample. List of cytokines included in the array is provided by a link in Table S5.

To further confirm protein expression, ELISA was used for GRO, CXCL8 and IL6. Briefly, TNBC cells were either treated or untreated with GST-TRAIL for 24hours in SFM with subsequent collection of the supernatant for ELISA.

For ELISA with mouse tumor tissue, tumors from control and APO-2L treated animals were collected, weighed, and homogenized (Table S5) supplemented with protease inhibitor cocktail according to manufacturer’s protocol to prepare the tumor lysates. Human ELISA kits were used for tumor lysates and serum isolated from murine samples (Table S5).

### Western Blot

Western blots were performed with TNBC cell lysate as previously described [14]. Murine tumor tissue lysates were prepared using tissue extraction reagent (Table S5) supplemented with protease inhibitor cocktail for western blotting. The antibodies used are listed in Table S5.

### Caspase-3/7 activity assay

Caspase 3/7 assays were performed as previously reported [14]. Caspase 3/7 activity was measured in donor isolated neutrophils incubated in SFM, SFM-T, CM, T-CM for the indicated time points.

### Chromatin immunoprecipitation assay (ChIP)

MDA-MB231 cells were cultured with or without TRAIL for 2h before. ChIP assays were performed by the Magnetic ChIP Kit (Table S5) according to manufacturer’s instructions. DNA-bound protein was immunoprecipitated using anti-p100/p52 antibody or rabbit IgG, as a control. The eluted DNA was quantified by real-time PCR using specific primer sets (Table S5) flanking the expected promoter site of each gene [77].

### Immunofluorescence microscopy of neutrophils

Polarized morphology of neutrophils and actin expression were studied by immunofluorescence microscopy. Briefly, 18mm coverslip were pre coated with 25ug/ml of fibrinogen dissolved in DPBS for 30 min at 37℃. 5x10^5^ neutrophils were seeded on these coverslips for 10 min at 37℃ allowing them to settle on the coverslip. Cells were then stimulated with DMSO in SFM, fMLP (100nM). CM, or T-CM for 30 min at 37℃. Following stimulation, the cells were fixed with 4% formaldehyde for 15 min at room temperature (RT). Once fixed, the cells were permeabilized with 0.05% saponin in 3% BSA-containing DPBS for 15 min at RT and stained with phalloidin-FITC over night at 4℃. The cells were then stained and mounted with DAPI and imaged using Nikon Eclipse Ti2 / Yokogawa CSU-W1 confocal microscope with the 60X objective. Cells in at least 5 different fields were captured randomly per condition in each experiment. F-actin intensity was measured with image j software.

### Trans-well migration

To quantify the ability of conditioned media to induce neutrophil migration the Cytoselect^TM^ 24- well cell migration assay (Table S5) was used according to manufacturer’s protocol. Briefly, freshly isolated neutrophils suspended in SFM were placed inside each insert, and migration was observed for 90 min against different breast cancer cell-derived CM or T-CM. To assess the role of TRAIL on migration of neutrophils (exogenous GST-TRAIL 45nM) was added to the SFM or CM. To investigate the impact of NFKB1 and NFKB2 gene on TRAIL-mediated neutrophil migration, trans-well migration of freshly isolated neutrophils was performed against supernatant from TNBC cells transfected with NFKB1 and NFKB2 siRNA. For inhibition studies neutrophils were either incubated overnight with the CXCR1/2 inhibitor 1uM Reparixin (1 μM) or the IL6RA inhibitor Sarilumab (200 μg/ml), followed by chemotaxis assays against SFM, CM or T-CM as described above. To investigate the effect of neutrophil secreted factors in response to TRAIL on other neutrophils, migration of freshly isolated neutrophils was observed against ‘neutrophil condition media’.

To quantify the ability of murine conditioned media to induce murine neutrophil migration similar protocol was followed as above. Neutrophils were isolated from murine whole blood collected by cardiac puncture in K2 EDTA tubes using kits and reagent mentioned in Table S5.

### In vivo model

All experiments using animals were performed on approved studies in accordance with the guidelines provided by the National Cancer Institute (National Institutes of Health, Bethesda, MD, USA) Animal Care and Use Committee (ACUC) and were compliant with all relevant ethical regulations regarding animal research. In all experiments, female 5-week-old athymic NRC nu/nu mice were used.

In the first set of experiment, we investigated the TRAIL induced cytokine production by tumors. 5 × 10^6^ MDA-MB-231T human breast cancer cells were implanted into the mammary fat pad (MFP) of mice. The tumor size was monitored weekly by caliper measurements (e.g., tumor size [mm^3^] = length [mm] × width[mm] ^2^× 0.52). Once the average tumor size reached 200 mm^3^, mice were randomized into treatment and control groups (10 mice/group). Recombinant TRAIL/Apo2L (100 ug/200ul/ body weight) or control vehicle (PBS) was administered via tail vein injection, The mice were sacrificed 24hours post treatment, the tumors were dissected, measured, a section was stored in RNAlater (for RNA isolation), and protein lysates was prepared using tissue extraction reagent (Table S5) supplemented with protease inhibitor cocktail. Whole blood was collected in K2 EDTA tubes and serum was collected after resting it for 30 min at room temperature followed by centrifugation (20000g for 10 min at 4℃). Serum was stored at -80℃ storage until cytokine measurement.

### CODEX and immunohistochemistry (IHC) or Myeloperoxidase (MPO)

To investigate the recruitment of neutrophils in response to TRAIL, fixed frozen tissue from our previous study with the DR5 agonist MEDI3039 [14] was analyzed with Co-detection by indexing (CODEX) and IHC of myeloperoxidase (MPO) in tumor tissue. CODEX was performed as described previously [78]. Antibodies used for CODEX are listed in Table S5.

IHC of MPO was performed using 5-micron paraffin embedded sections on a Leica Bond RX auto stainer (Leica Biosystems). MPO primary antibody was diluted to 1:500, with a 30- minute incubation time. Antibody detection was accomplished using the Bond Polymer Refine Detection kit (Leica Biosystems) with the post-primary reagent removed from Leica’s default staining protocol. Myeloid leukemia mouse liver tissue was used as a positive control.

### Intra vital Microscopy

The recruitment of neutrophils to the tumor microenvironment in response to TRAIL was also observed using intravital microscopy. 3x10^5^ Dendra2 labelled MDA- MB- 231 cells were injected in the mammary fat pad (MFP) of 6-8 weeks athymic nude female mice, 4 mice per arm. 4 weeks after tumor development, mice were injected with APO 2L (100 μg/200ul/ body weight) i.v. via the tail vein. 24hours post TRAIL injection, neutrophils from the bone marrow of normal athymic nude mice were isolated and labelled with Cell Tracker as previously described [79]. 5 x10^6^ labelled neutrophils were injected into either the APO2L or saline-treated tumor bearing mice. Intravital microscopy was performed as previously described beginning 6hours post neutrophil injection till 4h [80]. Briefly, the mice were anesthetized using 1% isoflurane followed by an intraperitoneal injection of a mixture of 100 mg/kg and 10 mg/kg xylazine) in 0.9% sterile saline solution. The mice were placed on a heating pad throughout the procedure to maintain the body temperature at 37°C. The fur was shaved from the skin surrounding the mammary gland tumor, and an incision was made by pinching the skin with tweezers and cutting the raised skin with blunt-ended scissors. The exposed tumor was kept moist by applying carbomer 940 gel [80]. The mouse was then transferred to the heated stage and a custom-made spacer was used to stabilize the tumor. Imaging was performed by using an inverted TCS SP8 Dive Spectral Microscope equipped with a Mai-Tai laser (Spectral Physics), 3 spectral detectors (HyD-RLD, Leica) and a 37°C preheated 40X objective (NA 1.10, HC PL IRAPO, Leica). The specimens were excited at 930 nm. Collagen I (Second Harmonic Generation), Dendra2, and Cell Tracker Red were detected respectively in the following emission ranges: 461-471 nm, 494-528 nm, and 598-616 nm. Images were acquired in tile mode by bidirectional line scanning at 400 Hz (512 x 512 pixel; 12 bits per pixel) on XY and stitched using the LAS X Navigator software. The 3-D image of the recruitment of stained neutrophils were acquired by bidirectional line scanning with 3 lines averaging at 400 Hz (512 x 512 pixel; 12 bits per pixel) on XY, and 45 to 55 Z-stacks (2 um step size) using Leica LAS X software. All the images were stored as LIF files and processed further using Imaris software.

### Cytotoxicity assay

5000 luciferase labelled LM2-MDA-MB-231 cells were seeded in a flat bottomed 96 well plate in complete DMEM and incubated at 37℃ overnight. Donor isolated purified neutrophils were incubated in either CM or T-CM collected from TNBC cell lines for 16-18h at 37℃. Following the incubation, the neutrophils were centrifuged, condition media was aspirated, neutrophil pellets were washed with PBS, then cocultured with the target, MDA-MB-231 cells in SFM for 4h (≥12 replicates per condition) at target: effector ratios of 1:0, 1:1, 1:3, and 1:10. Following co-culture, the wells were washed with PBS and cancer cell viability was measured by luciferase activity using the luciferase reporter assay system (Table S5). During the 4hours incubation period, the viability of the neutrophils was confirmed by AOPI assay. Cytotoxicity percentage was calculated as [1- (luciferase activity in cells incubated with neutrophils/ luciferase activity in target cells incubated without neutrophils)] x 100.

### Annexin V assay

The Annexin V Apoptosis and Necrosis Assay Kit (Table S5) was used to detect binding of Annexin V to neutrophil membranes under different conditions as indicated in the text according to the manufacturer’s protocol.

### T cell proliferation assay

To observe the effect of T-CM incubated neutrophils on T cell proliferation, PBMC isolated from healthy blood donors were stained with CFSE and (5 × 10^4^) seeded in a 96 well round bottom plate followed by activation using CD3/CD28 dyna beads. This was followed by co-culturing with autologous neutrophils (1:1 ratio) preincubated in different condition media (SFM, SFM-T, CM, T-CM). The co culture was performed in complete RPMI media for 72 hours followed by staining with antibodies listed in Table S5 for flow cytometry analysis. FlowJo software was used to evaluate the CFSE dilution rate of T cells.

### Statistical analysis

Graph Pad Prism 10.1.1 software was used for the statistical analyses. All quantitative data are presented as the mean ± standard deviation of error (SEM) or mean ± standard deviation (SD). The significance of differences in data was determined using either Student’s *t*-test, or 2-way ANOVA as indicated in figure legends.

## Acknowledgement

We thank Dr. Jan Wisniewski (Experimental Immunology Branch, National Cancer Institute, Bethesda, USA), Dr. Elijah Edmondson (Molecular Histopathology Lab, Laboratory of Animal Sciences Program, CCR, NCI, Frederick), Angie Schwab (Center for Immuno-Oncology, CCR, NCI) for technical assistance with immune assays, Abigail J. Walke (Optical Microscopy and Analysis Laboratory, Cancer Research Technology Program, Frederick National Laboratory for Cancer Research, Frederick, MD 21702, USA.) for helping in CODEX imaging, and Dr. Massimo Fantini (Women’s Malignancies Branch, CCR, NCI), for technical assistance of flow cytometry experiments. We also would like to thank members of the Lipkowitz lab for their discussion and support.

## Author Contributions

M. Kundu: Conceptualization, data curation, software, formal analysis, validation, investigation, visualization, methodology, writing-original draft, writing-review and editing. Y.E. Greer: Investigation, visualization, data curation, writing-review and editing. A. Lobanov. Investigation, analysis, data curation and editing. L. Ridnour, R.N. Donahue, Y. Ng. and S. Ratnayake: Investigation, data curation, analysis and editing. D. Voeller and S. Weltz: Investigation, writing-review and editing. Q. Chen: data curation and editing. S.J. Lockett: methodology. M. Cam, D. Meerzaman, D. A. Wink, and R. Weigert: Resources, data curation, supervision and editing. S. Lipkowitz: Resources, formal analysis, supervision, funding acquisition, project administration, writing-review and editing.

## Ethics Approval

All experiments using animals were performed on approved studies in accordance with the guidelines provided by the National Cancer Institute (National Institutes of Health, Bethesda, MD, USA) Animal Care and Use Committee (ACUC) and were compliant with all relevant ethical regulations regarding animal research. Human peripheral blood was obtained from healthy donors on an IRB-approved NIH protocol (99-CC-0168). Research blood donors were at least 18 years of age and provided written informed consent. All blood samples were de-identified prior to distribution. Clinical Trials Number: NCT00001846. All experimental procedures were performed in accordance with the recognized ethical guidelines of the Declaration of Helsinki.

