## Supplemental information for "TRAIL-induced cytokine production via NFKB2 pathway promotes neutrophil chemotaxis and immune suppression in triple negative breast cancers"

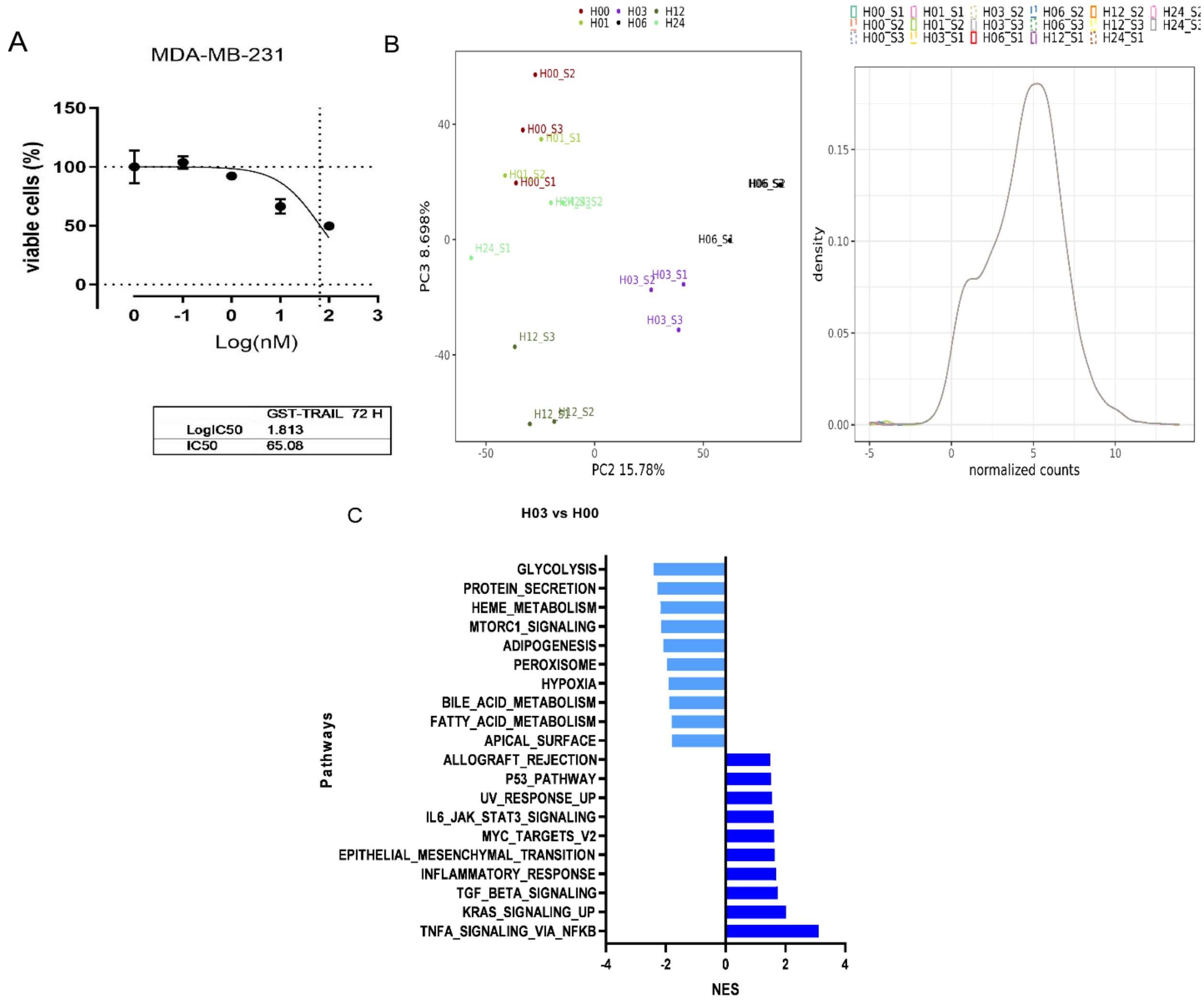

**Supplementary Figure 1. TRAIL treatment causes transcriptional change in MDA-MB-231 cells.** A. MTS assay in MDA-MB-231 cells treated with GST-TRAIL for 72h. Calculated IC50 was 65.08. Data shown is the mean +/-SEM of 3 independent experiments. B. Left panel- Principal component analysis (PCA) plot of MDA-MB-231 cells treated with 45 nM of TRAIL at different time points as indicated. Right panel- Density plot of normalized data from RNA seq of MDA-MB-231-time course in response to TRAIL treatment using Relative Log Expression (RLE) normalization method based on read count and log2. C. GSEA for the hallmark set of genes showing pathways enriched at the 3h time point. The plot shows the top 10 enriched pathways (dark blue) and top 10 decreased pathways (light blue) at 3h post TRAIL treatment in MDA-MB-231 cells.

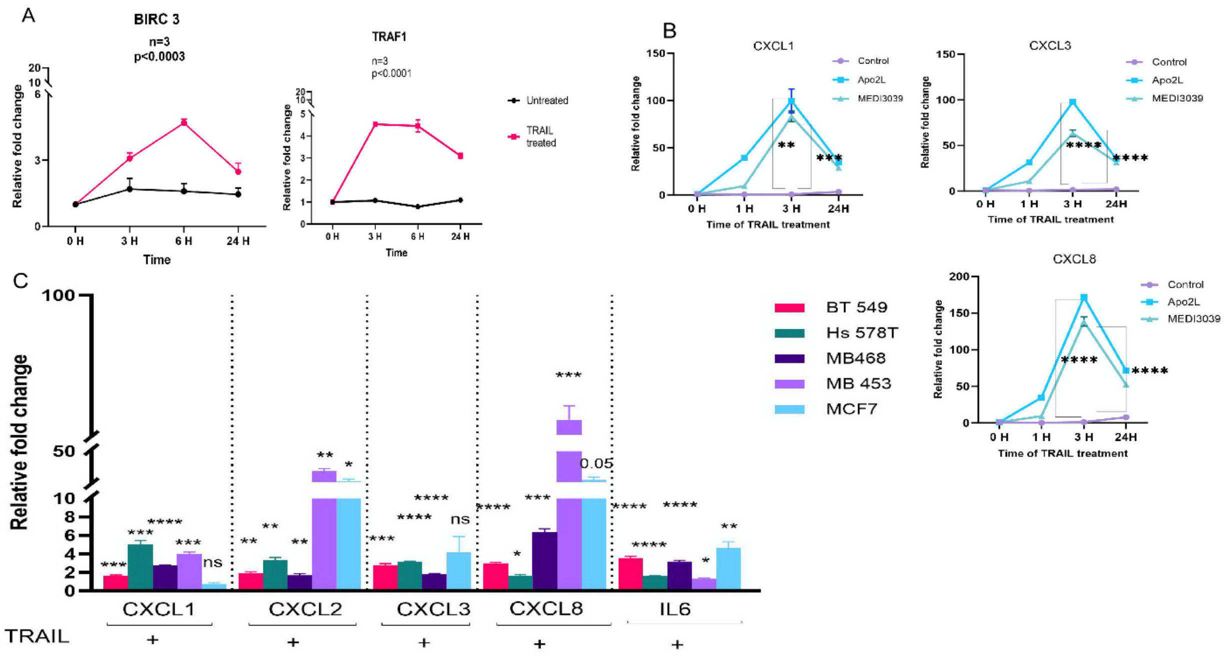

**Supplementary Figure 2. TRAIL induces the expression of immunomodulatory genes in TNBC cells** A. Time course qRT-PCR of BIRC 3 and TRAF1 with TRAIL treatment in MDA-MB-231 cells and the data are the mean +/-SEM of 3 independent experiments; 2-way ANOVA was used for the comparison. B. TRAIL agonist MEDI3039 and Apo2L induced cytokine transcripts in MDA-MB-231T cells at 3h time point measured by qRT-PCR. 2-way ANOVA was used for all the time points and student t- test for 3h time point. Data are the mean +/-SEM of 3 independent experiments C. TRAIL induced cytokines mRNA measured by qRT-PCR (CXCL1, CXCL2, CXCL3, CXCL8, and IL6) in the indicated breast cancer cells. Data are the mean +/-SEM of at least 3 independent experiments. *$p$<0.05, **$p$<0.01, ***$p$<0.001, ****$p$<0.0001.

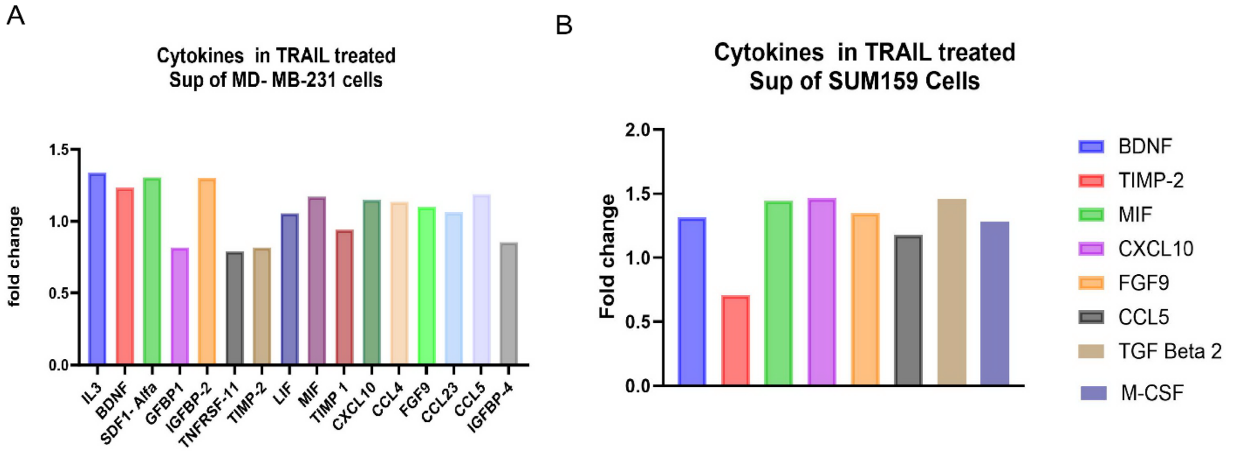

**Supplementary Figure 3. Cytokine array showing altered expression of proteins in response to TRAIL** A. Quantification of cytokine array showing the expression of various cytokines in TRAIL treated vs untreated supernatants from MDA-MB-231 cells. H. Cytokine array showing the expression of various cytokines in TRAIL treated vs untreated supernatant of SUM159. Data was normalized to positive control according to manufacturer's protocol.

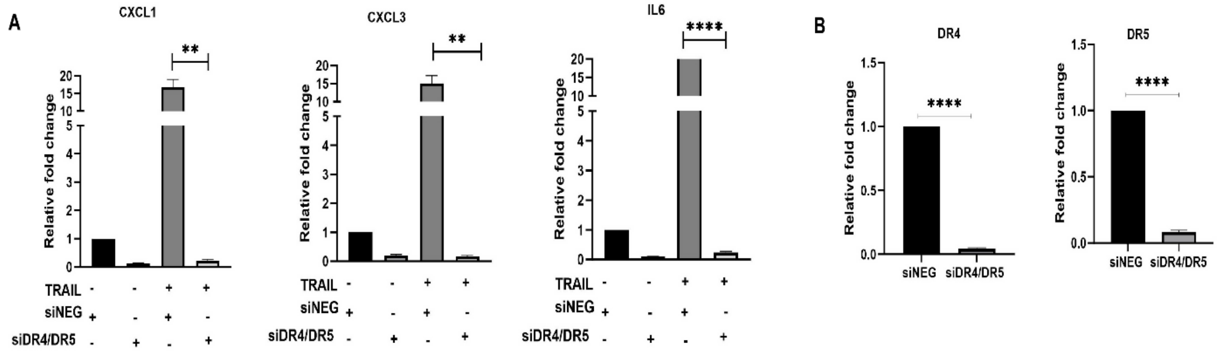

**Supplementary Figure 4: TRAIL induces cytokine mRNA via DR4/DR5.** A. qRT-PCR data showing effect of simultaneous DR4/DR5 siRNA-mediated knockdown of DR4 and DR5 on representative TRAIL induced cytokines. B. Confirmation of the knock down of both DR4 and DR5 in MDA-MB-231 cells by qRT-PCR. Data are the mean +/-SEM of 3 independent experiments. ****p*<0.01, ******p*<0.0001, Student's *t*-test.

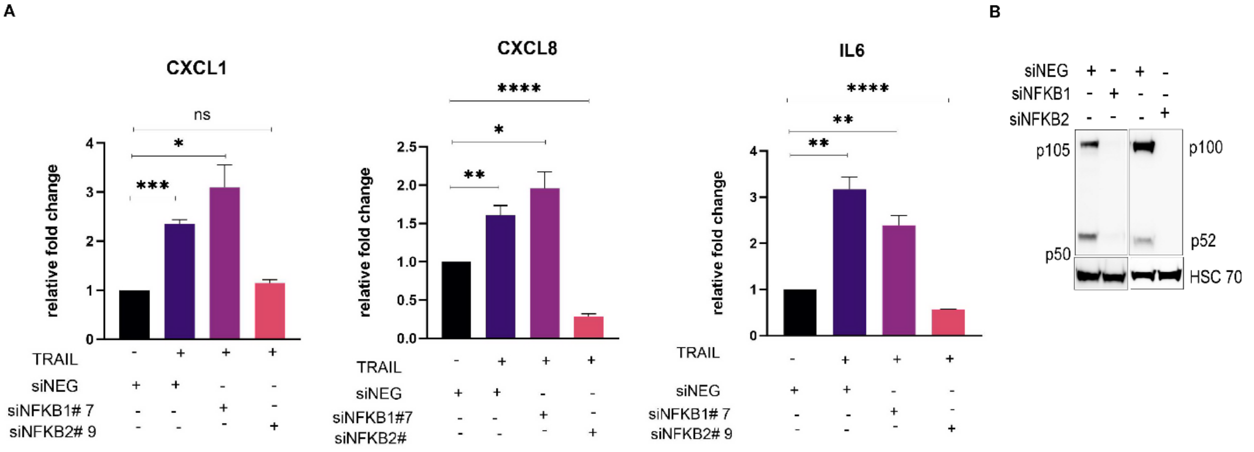

**Supplementary Figure 5: Compared to NFKB1, NFKB2 knockdown had greater effect on TRAIL-induced cytokine production**. **A.** TRAIL-induced cytokine mRNA level in MB-MDA-231 cells with siRNA mediated knockdown of NFKB1 or NFKB2 using qRT-PCR at 3h time point post GST-TRAIL treatment using additional siRNAs (compared to those used in Fig. 4 D and F). Data are the mean +/- SEM of 3 independent experiments. **p*<0.05, ***p*<0.01, ****p*<0.001, *****p*<0.0001, Student's *t*-test. B. Immunoblot confirming the knock down of NFKB1 and NFKB2 protein compared to siNEG.

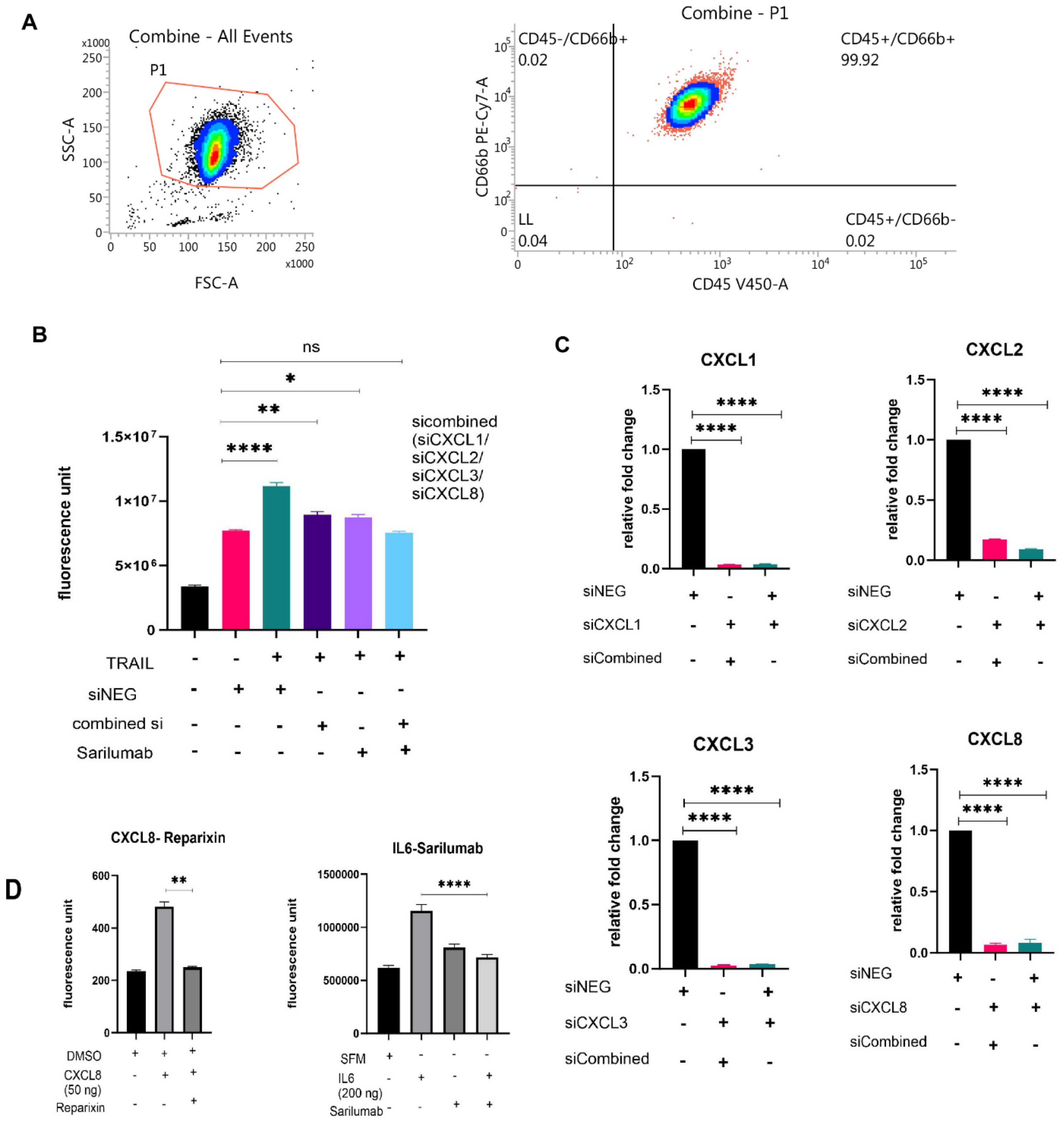

**Supplementary Figure 6. A. Characterization of neutrophils isolated from whole blood of healthy donors by using negative selection kit (Table S5).** Isolated neutrophils were stained with CD45, CD66B and live dead stain and FLOW cytometry was performed (representative assay shown). Isolated neutrophils used in assays described in main text were analyzed by Flow jo. Live

CD45+CD66B+ cells were characterized as human neutrophils; the assay demonstrated 99.9% of isolated cells wereCD45+CD66B+ cells indicating purified population of isolated neutrophils.

**B. Combination of cytokines (CXCL1, 2, 3, 8 and IL6) in T-CM are responsible for TRAIL induced neutrophil chemotaxis.** Chemotaxis of neutrophils against T-CM collected from MDA-MB-231 treated +/- GST-TRAIL after siRNA mediated combined knockdown of the cytokines (CXCL1/CXCL2/CXCL3/CXCL8) or siNEG. The chemotaxis assays were conducted with the T-CM from the knockdown cells +/- the IL6 receptor inhibitor Sarilumab. C. Confirmation of knockdown of the mRNA for each indicated gene i by RT-PCR. D. Chemotaxis of human isolated neutrophils against MDA-MB-231 CM supplemented with recombinant human CXCL8 or IL6 to confirm the effect of Reparixin (CXCR1/CXCR2 blocker and Sarilumab (IL6Receptor inhibitor).

Data represented in B-D are the mean+/- SEM of 3 independent experiments; Student's *t*-test was used *$p<0.05$, **$p<0.01$, ****$p<0.0001$.

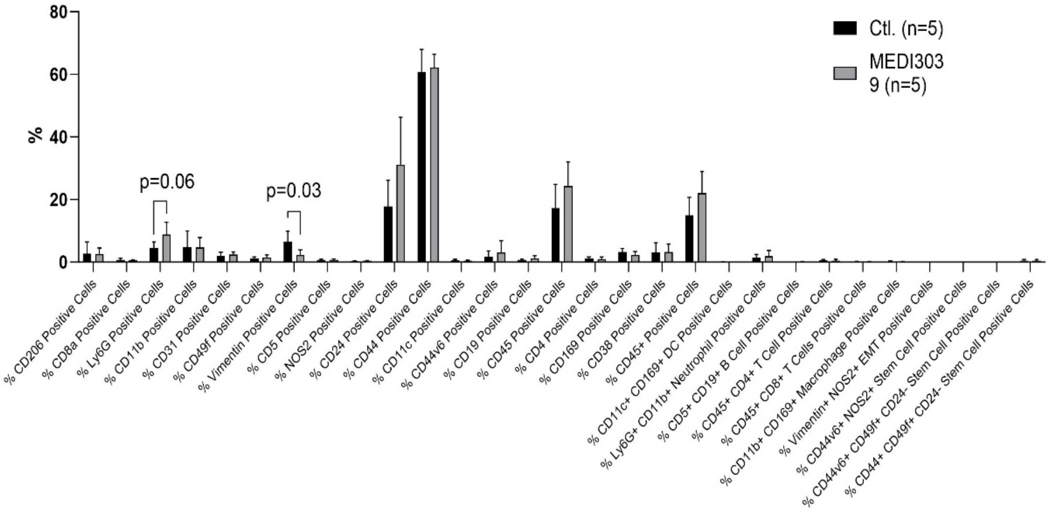

**Supplementary Figure 7. TRAIL agonist MEDI3039 promotes neutrophil recruitment in the tumor tissue as analyzed by CODEX.** Graphical representation of CODEX analysis showing percentage of various immune cells in tumor tissue from mice treated with MEDI3039 (n=5) compared to those treated with vehicle control (n=5). Human vimentin+ cells indicate presence of MDA-MB-231 TNBC cells. Data shown is the mean+/- SEM.

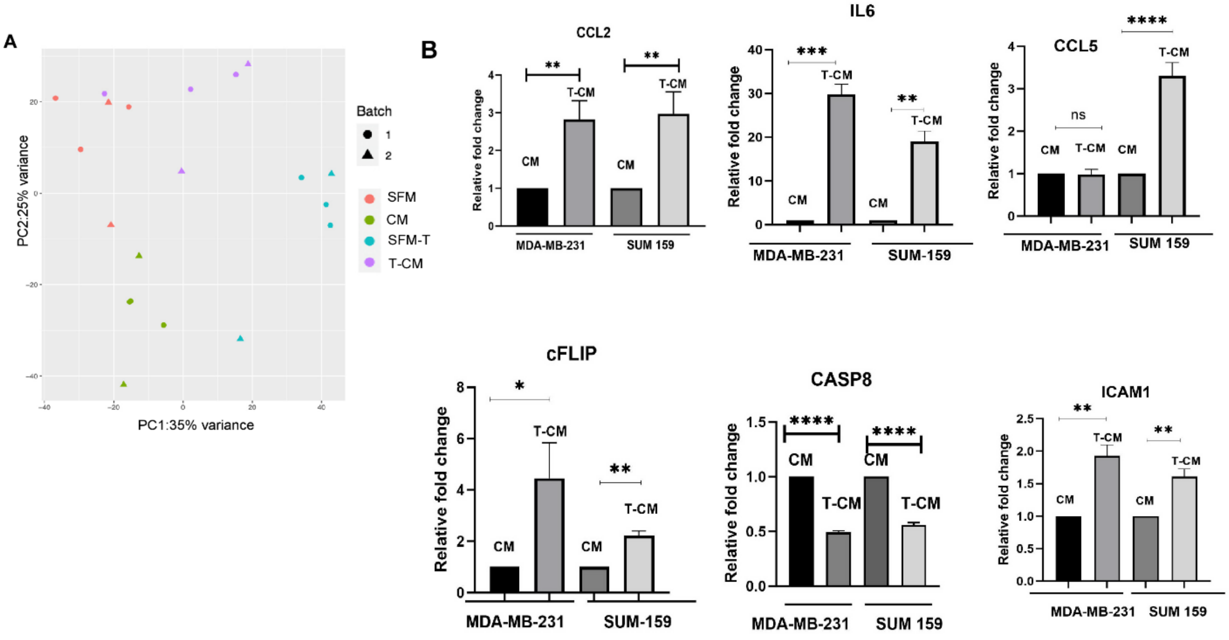

**Supplementary Figure 8. T-CM and TRAIL increase the expression of immunomodulatory genes in the neutrophils. A.** Principal component analysis (PCA) plot of neutrophils incubated in SFM/SFM-T/CM/T-CM based on RNAseq data. B. qRT-PCR Expression of immunoregulator and apoptosis gene mRNAs measured by qRT-PCR in neutrophils in response neutrophil incubation in T-CM from TNBC cells (MDA-MB-231 and SUM159). Data are the mean +/-SEM of multiple independent experiments (n-3-5).

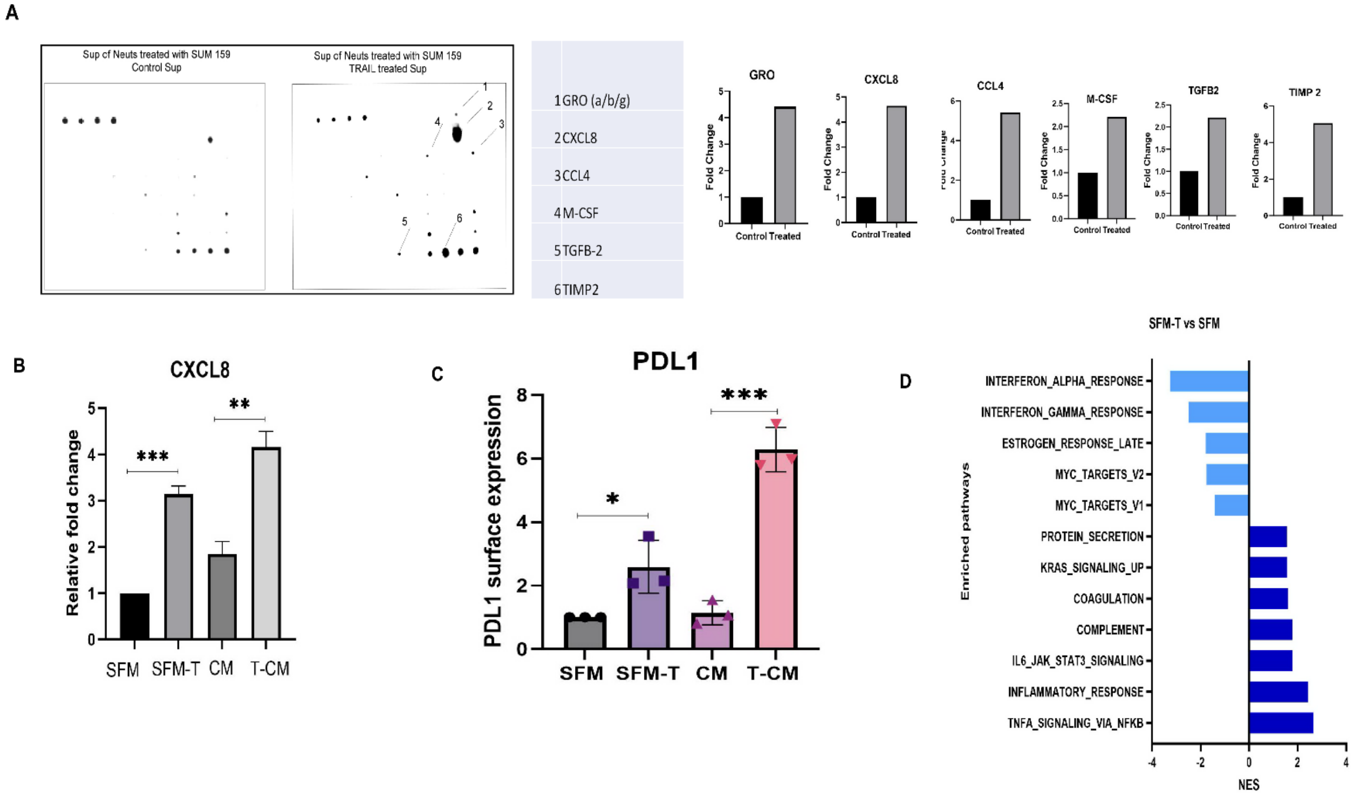

**Supplementary Figure 9. T-CM and TRAIL increase the protein expression of immunomodulatory genes in the neutrophils.** A. Cytokine protein expression in neutrophils incubated in SFM vs T-CM from SUM159 measured by protein array (top) and quantitated (below) s. B. Upregulation of CXCL8 mRNA in response to TRAIL and T-CM from MDA-MB-231 cells (the upregulation was identified in the cytokine array and not in the RNAseq). Data are the mean +/-SEM of 3 independent experiments. C. Surface expression of PDL1 by FLOW cytometry on neutrophils isolated from healthy donor blood incubated in SFM, SFM-T, CM and T-CM from MDA-MB-231 cells. Data are the mean +/-SD of n=3 independent experiments. D. GSEA for the hallmark set of genes from neutrophils incubated in SFM-T vs SFM (upregulated pathways-dark blue; down regulated pathways-light blue). *$p<0.05$, **$p<0.01$, ***$p<0.001$, ****$p<0.0001$, Student's $t$-test.

**A**

| Murine cell names | Subtypes | M-TRAIL sensitivity | Fold of mRNA change | Chemotaxis assay |
|---|---|---|---|---|
| 4T1 | Epithelial TNBC | IC50> 1000nM | 4 | N.D. |
| 4T1.2 | Single cell clone of 4T1 | IC50> 100nM | 2.5 | N.D. |
| MET 1 | TNBC | IC50> 1000nM | 40 | Increased with >1000nM of TRAIL |
| AT3 | Autochthonous Tumor (TNBC) | IC50> 1000nM | 6 | Increased with >1000nM of TRAIL |
| M6 | TNBC | Not performed | 2 | |
| MVT1 | Mesenchymal TNBC | IC50> 1000nM | Not tested | Not tested |
| E0071 | Mesenchymal TNBC | IC50> 1000nM | Not tested | Not tested |

**B**

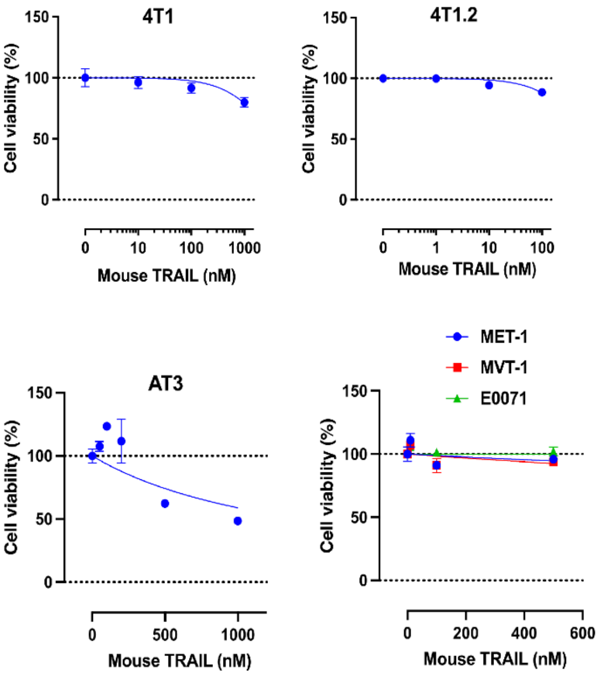

**Supplementary Figure 10. Effect of TRAIL on murine TNBC cell lines. A.** Summary table showing the effect of mouse TRAIL (M-TRAIL) on murine TNBC cell lines. N.D. (Not Detected). B. MTS assay over 72h showing the percentage of viable cells in response to indicated concentration of M-TRAIL in murine TNBC cell lines. Data are the mean +/-SEM of 3 independent experiments.

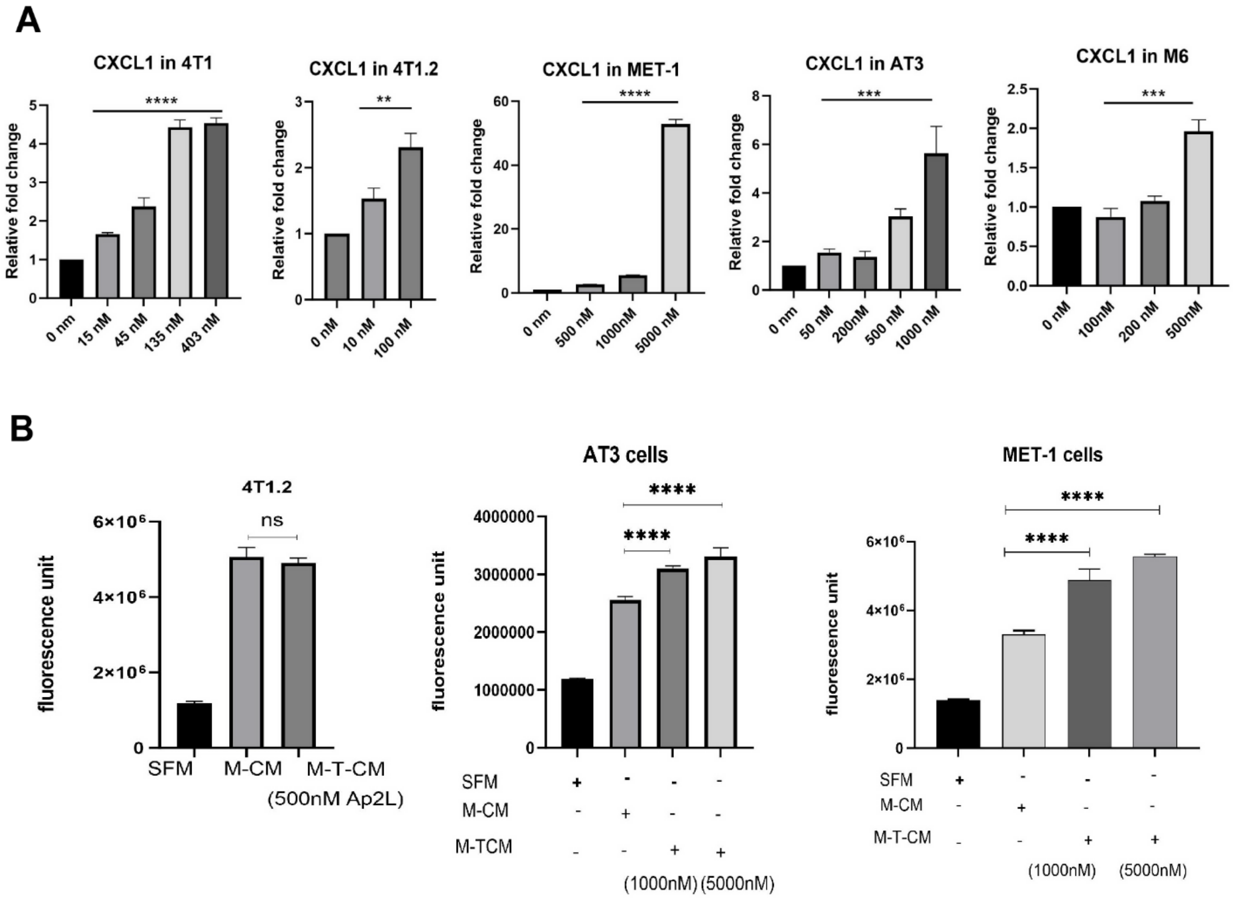

**Supplementary Figure 11. Effect of TRAIL on murine cytokine expression and murine neutrophil chemotaxis. A**. TRAIL induced cytokine expression (CXCL1) in murine TNBC cells in response to indicated dose of human Apo2L. Data are the mean +/-SEM of 3 independent experiments. One way ANOVA of multiple comparison with control was used. B. Chemotaxis assay of murine neutrophils in response to mouse TRAIL- treated Condition media (M-T-CM) and mouse -tumor condition media (M-CM). Data is as the mean +/-SEM of 3 independent experiment using Student's *t*-test. ***p*<0.01, ****p*<0.001, *****p*<0.0001.

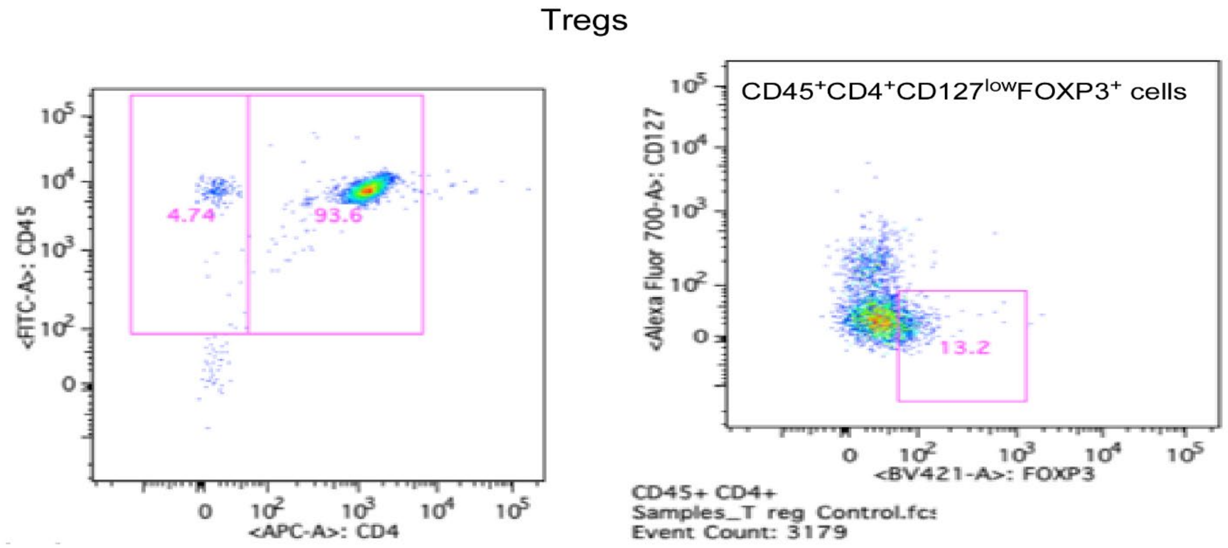

**Supplementary Figure 12. Characterization of Tregs isolated from whole blood of healthy donors by FLOW cytometry.** Treg cells isolated following manufacturer's protocol were stained with CD45, CD4, CD127 and FOXP3 antibodies. The data was further analyzed by Flow jo and gated for CD45+CD4+CD127 lowFOXP3+ cells which were characterized as Treg cells.

**Table S2: The top 25 differentially up-regulated genes at 3h and 24h in MDA-MB-231 cells in response to TRAIL**

| 3H vs 0H | | 24H vs 0H | |
|---|---|---|---|
| Gene Name | logFC | Gene Name | logFC |
| CXCL1 | 2.9 | AC131160.1 | 6.3 |
| MIR3142HG | 2.7 | UNC13A | 3.2 |
| CXCL8 | 2.4 | CXCL11 | 2.9 |
| CSF2 | 2.4 | CTSW | 2.1 |
| BIRC3 | 2.2 | VCAM1 | 2.1 |
| NUAK2 | 2.0 | BIRC3 | 2.0 |
| LTB | 1.8 | TNFRSF9 | 1.8 |
| JUNB | 1.7 | CSF3 | 1.8 |
| IL6 | 1.7 | AP000781.2 | 1.7 |
| CXCL3 | 1.7 | SSTR2 | 1.6 |
| TNFAIP3 | 1.6 | TRAF1 | 1.6 |
| PDGFB | 1.6 | TNFSF15 | 1.6 |
| TRAF1 | 1.5 | PDK4 | 1.5 |
| NR4A1 | 1.5 | CXCL1 | 1.5 |
| IL11 | 1.5 | GBP4 | 1.4 |
| AC092807.3 | 1.5 | GPRC5B | 1.3 |
| CXCL2 | 1.4 | ICAM1 | 1.3 |
| NFKBIA | 1.4 | PECAM1 | 1.3 |
| ZC3H12A | 1.4 | CNGA1 | 1.3 |
| KRTAP2-3 | 1.3 | EGF | 1.3 |
| NFKBIE | 1.3 | BACH2 | 1.2 |
| CYP1A1 | 1.2 | NGFR | 1.2 |
| RELB | 1.2 | NR3C2 | 1.2 |
| AL138724.1 | 1.2 | KYNU | 1.2 |
| AC020916.1 | 1.2 | PDE4B | 1.2 |

**Table S3**

| Breast Cancer Cell Types | Subtypes | | | Culture media | Serum |
|---|---|---|---|---|---|
| ***Human*** | ***ER*** | ***PR*** | ***HER2*** | | |
| MDA-MB-231 | - | - | - | RPMI | 10%FBS |
| MDA-MB-231T | - | - | - | DMEM | 10%FBS |
| LM2-MDA-MB-231 | - | - | - | DMEM | 10%FBS |
| Dendra2-MDA-MB-231 | - | - | - | DMEM | 10%FBS |
| SUM 159 | - | - | - | DMEM F12* | 5% FBS |
| BT 549 | - | - | - | RPMI | 10%FBS |
| Hs578T | - | - | - | RPMI | 10%FBS |
| MDA-MB-468 | - | - | - | RPMI | 10%FBS |
| MDA-MB-453 | - | - | + | RPMI | 10%FBS |
| MCF-7 | + | - | - | RPMI | 10%FBS |
| **Murine** | | | | | |
| **4T1** | - | - | - | DMEM | 10%FBS |
| **4T1.2** | - | - | - | RPMI | 10%FBS |
| **MET-1** | - | - | - | DMEM | 10%FBS |
| **MVT-1** | - | - | - | DMEM | 10% FBS |
| **E0071** | - | - | - | RPMI** | 5% FBS |
| **M6** | - | | | DMEM | 5%FBS |
| **AT3** | + | - | - | DMEM | 10%FBS |

### Table S3 Legend:

ER, estrogen receptor; PR, progesterone receptor; HER2, human epidermal growth factor receptor 2. Culture media conditions: FBS, fetal bovine serum supplemented with 100 units/ml of penicillin, 100 ug/ml of streptomycin. *Media supplemented with 5ug/ml insulin, 1ug/ml hydrocortisone. **Media supplemented with 10 mmol/L HEPES. Dendra-2 MDA- MB-231 and LM2 MDA MB-231 were gifts from Dr. Roberto Weigert and Dr. Esta Sterneck, NCI/CCR, respectively. SUM 159 was a gift from Dr. Lee Graves, UNC and the murine cell lines were provided by Dr. Lalage Wakefield, NCI, Bethesda, MD. The other cell lines were purchased from American Type Culture Collection (ATCC).

**Table S5: Reagents**

**Chemicals, reagents, assay kits**

| Reagent/Kit | Source | Identifier/catalog number |
|---|---|---|
| Cellometer ViaStain™ AOPI Staining Solution | Nexcelcom Bioscience, Lawrence, MA | CS2-0106-5mL |
| CellTiter-Glo® 2.0 Cell Viability Assay | Promega, Madison, WI | G9242 |
| Sarilumab | SelleckChem | A2011 |
| Reparixin | Sigma-Aldrich, St. Louis, MO | SML2655 |
| Human IL2 | STEMCELL Technologies, Inc., Vancouver, Canada | 78036 |
| Dimethylsulfoxide (DMSO) | Sigma-Aldrich, St. Louis, MO | D2650-100ml |
| Recombinant Human CXCL8 | R and D Systems, Minneapolis, MN | 208-IL-010 |
| Recombinant Human IL6 | R and D Systems, Minneapolis, MN | 206-IL/CF |
| Lipofectamine® RNAiMax | ThermoFisher Scientific, Waltham, MA | 13778150 |
| Bright-Glo™ EX Luciferase Assay System | Promega, Madison, WI | E2610 |
| Caspase-Glo® 3/7 Assay System | Promega, Madison, WI | G8090 |
| RealTime-Glo™ Annexin V Apoptosis and Necrosis Assay Kit | Promega, Madison, WI | JA1011 |
| Apo 2L (Human) | Promega, Madison, WI | 310-04 |
| SuperKillerTRAIL™, Soluble (mouse) | Adipogen Lifesciences, San Diego, CA | AG-40T-0004 |
| Glycogen, RNA grade | ThermoFisher Scientific, Waltham, MA | R0551 |
| Pierce Magnetic ChIP kit | ThermoFisher Scientific, Waltham, MA | 26157 |
| 16%Formaldehyde | Cell Signaling, Danvers, MA | 12606P |
| Fibrinogen from human plasma | Sigma-Aldrich, St. Louis, MO | F4883 |
| N-Formyl-Met-Leu-Phe (fMLP) | Sigma-Aldrich, St. Louis, MO | F3506 |
| Saponin | Sigma-Aldrich, St. Louis, MO | 84510 |
| BSA | Sigma-Aldrich, St. Louis, MO | A3294-500G |
| Ammonium Chloride Solution | STEMCELL Technologies, Inc., Vancouver, Canada | 7800 |
| Duolink® *In Situ* Mounting Medium with DAPI | Sigma-Aldrich, St. Louis, MO | DUO82040 |
| Cytoselect™ 24-well cell migration assay | Cell Biolabs, INC., San Diego, CA | CBA-103-5 |
| RayBio® C-Series Human Cytokine Antibody Array C5 (List of Cytokines) | RayBiotech, GA | https://www.raybiotech.com/human-cytok |
| RayBio® C-Series Human Cytokine Antibody Array C5 | RayBiotech, GA | AAH-CYT-5-4 |
| Human GRO ELISA Kit | RayBiotech, GA | ELH-GRO |
| Human IL-8/CXCL8 ELISA Kit - Quantikine | R and D Systems, Minneapolis, MN | D8000C |
| Human IL-6 Quantikine ELISA Kit | R and D Systems, Minneapolis, MN | D6050 |
| Tissue extraction Reagent 1 | ThermoFisher Scientific, Waltham, MA | FNN0071 |
| Human ELISA Kit CXCL8 (Tissue lysates) | RayBiotech, GA | ELH-IL8-CL |
| Human ELISA Kit IL6 (Tissue lysates) | RayBiotech, GA | ELH-IL6-CL |
| Human ELISA Kit CXCL8 (serum) | RayBiotech, GA | ELH-IL8 |
| Human ELISA Kit IL6 (serum) | RayBiotech, GA | ELH-IL6 |

**Materials for molecular biology experiments**

| Reagent/Kit | Source | Identifier/catalog number |
|---|---|---|
| RNeasy Mini Kit (50) | Qiagen, Germantown, MD | 74104 |
| TRIZOL | ThermoFisher Scientific, Waltham, MA | 15596018 |
| Power Up SYBR Green Master Mix | ThermoFisher Scientific, Waltham, MA | A25780 |
| QuantiTect Reverse Transcription Kit | Qiagen, Germantown, MD | 205313 |
| Rnase-Free Dnase Set | Qiagen | 79254 |

| Primer for qPCR | Source | Identifier/catalog number |
|---|---|---|
| Hs_CXCL1_1_SG | Qiagen | QT00199752 |
| Hs_CXCL2_1_SG | Qiagen | QT00013104 |
| Hs_CXCL3_1_SG | Qiagen | QT00015442 |
| Hs_CXCL8_1_SG | Qiagen | QT00000322 |
| Hs_CXCL11_2_SG | Qiagen | QT02394644 |
| Hs_IL6_1_SG | Qiagen | QT00083720 |
| Hs_BIRC3_1_SG | Qiagen | QT00021798 |
| Hs_TRAF_1_SG | Qiagen | QT00095732 |
| Hs_TNFRSF10A_1_SG (DR4) | Qiagen | QT00065723 |
| Hs_TNFRSF10B_1_SG (DR45) | Qiagen | QT00082768 |
| Hs_NFKB1_1_SG | Qiagen | QT00063791 |
| Hs_NFKB2_1_SG | Qiagen | QT00037590 |
| Hs_SIGLEC5_1_SG | Qiagen | QT00012404 |
| Hs_CD274_1_SG | Qiagen | QT00082775 |

| | | |
|---|---|---|
| Hs_CCL1_1_SG | Qiagen | QT00203154 |
| Hs_CCL2_1_SG | Qiagen | QT00212730 |
| Hs_CCL5_1_SG | Qiagen | QT00090083 |
| Hs_IL1A | IDT, Coralville, IA | 411360288 |
| Hs_IL1B | IDT, Coralville, IA | 420042959 |
| Hs_CFLAR_1_SG | Qiagen | QT00064554 |
| Hs_CASP8_1_SG | Qiagen | QT00052416 |
| Hs_ICAM1_1_SG | Qiagen | QT00074900 |
| Hs_TNF_1_SG | Qiagen | QT00029162 |
| Hs_ARG_1_SG | Qiagen | QT00068446 |
| Hs_FOXP3_1_SG | Qiagen | QT00048286 |
| Hs_CCR8_3_SG | Qiagen | QT02423834 |
| Hs_GAPDH_1_SG | Qiagen | QT00079247 |
| Hs_ACTB_1_SG | Qiagen | QT00095431 |
| Mm_Il6_1_SG | Qiagen | QT00098875 |
| Mm_Cxcl3_1_SG | Qiagen | QT00151599 |
| Mm_Cxcl1_1_SG | Qiagen | QT00115647 |
| Mm_Actb_1_SG | Qiagen | QT01136772 |
| siRNA | Source | Identifier/catalog number |
| all stars negative control siRNA | Qiagen | 1027281 |
| Hs_TNFRSF10A_1FlexiTube siRNA (siDR4) | Qiagen | SI00056728 |
| Hs_TNFRSF10B_1FlexiTube siRNA (siDR5) | Qiagen | SI03094063 |
| Hs_NFKB1_10 FlexiTube siRNA | Qiagen | SI02662618 |
| Hs_NFKB1_7 FlexiTube siRNA | Qiagen | SI02654932 |
| Hs_NFKB2_1 FlexiTube siRNA | Qiagen | SI00300965 |
| Hs_NFKB2_9 FlexiTube siRNA | Qiagen | SI04219852 |
| Hs_IL8_5 FlexiTube siRNA | Qiagen | SI02654827 |
| Hs_CXCL3_1 FlexiTube siRNA | Qiagen | SI00032662 |
| Hs_CXCL2_3 FlexiTube siRNA | Qiagen | SI00032648 |
| Hs_CXCL1_1 FlexiTube siRNA | Qiagen | SI00357280 |
| Hs_IL6_2 FlexiTube siRNA | Qiagen | SI00012579 |
| Hs_CASP8_12_GeneSolution siRNA | Qiagen | SI02662457 |
| Hs_CASP8_11_GeneSolution siRNA | Qiagen | SI02661946 |
| Hs_CASP8_7_GeneSolution siRNA | Qiagen | SI00299593 |
| Hs_CASP8_17_GeneSolution siRNA | Qiagen | SI04948314 |
| ChIP primers | | Identifier/catalog number |
| CXCL8 F:5_- GGGCCATCAGTTGCAAATC-3; | Eurofins, USA | 9024484 |
| R:5_GCTTGTGTGCTCTGCTGTCTC-3_ | | |
| CXCL2 F: 5_ATTCGGGGCAGAAAGAGAAC-3_ | Eurofins, USA | 9024484 |
| R:5_ ACCCCTTTTATGCATGGTTG-3 | | |
| GAPDH | ThermoFisher Scientific | 1862245 |
| ChIP Antibodies | | |
| p100/p52 antibody | Cell Signaling | D7A9K |
| Rabbit IgG | Cell Signaling | 2729 |
| **Reagents used for cell culture,Neutrophil, PBMC, Treg isolation kit** | | |
| Reagent/Kit | Source | Identifier/catalog number |
| DMEM, high glucose | ThermoFisher Scientific | 11965118 |
| DMEM/F12 medium | ThermoFisher Scientific | 11320082 |
| hydrocortisone | Sigma-Aldrich | H0888 |
| insulin | Sigma-Aldrich | I9278-5ML |
| RPMI 1640 medium | ThermoFisher Scientific | 11875119 |
| DPBS | ThermoFisher Scientific | 21-031-CV |
| LookOut® Mycoplasma PCR Detection Kit | Sigma-Aldrich | MP0035 |
| Promega GenePrint 10 System | Promega, Madison, WI | B9510 |
| Easy Sep™ Mouse Neutrophil Enrichment Kit | STEMCELL Technologies, Inc. | 19762 |
| Easy Sep™ Direct Human Neutrophil Isolation Kit | STEMCELL Technologies, Inc. | 19666 |
| Easy Sep™ Direct Human CD4+CD127lowCD25+ Regulatory T cell isolation Kit | STEMCELL Technologies, Inc. | 18063 |
| Dynabeads™ Human T-Activator CD3/CD28 for T Cell Expansion and Activation | ThermoFisher Scientific | 11131D |
| Ficoll-PaqueTM PLUS | GE Healthcare, Silver Spring, MD | 17-1440-02 |
| SepMate™-15 (IVD) | STEMCELL Technologies, Inc. | 85420 |
| **Materials for Western blotting** | | |
| Primary antibody | Source | Identifier/catalog number |
| NF-κB1 p105/p50 | Cell Signaling, Danvers, MA | 3035 |
| NF-κB2 p100/p52 | Cell Signaling, Danvers, MA | 4882 |

| | | |
|---|---|---|
| Phospho-NF-κB2 p100 (Ser866/870) | Cell Signaling, Danvers, MA | 4810 |
| DR4 | GeneTex, Irvine, CA | GTX28414 |
| DR5 | Santa Cruz Biotechnology | sc-65314 |
| Caspase-8 (1C12) Mouse mAb | Cell Signaling, Danvers, MA | 9746 |
| Cleaved- Caspase 8 | Cell Signaling, Danvers, MA | 9496 |
| Ly-6G (E6Z1T) Rabbit mAb | Cell Signaling, Danvers, MA | 87048 |
| Phospho-NF-κB p65 (Ser536) (93H1) Rabbit mAb | Cell Signaling, Danvers, MA | 3033 |
| beta Actin Antibody (C4) HRP | Santa Cruz Biotechnology, Dallas, TX | sc-47778 HRP |
| HSPA8/HSC70 Antibody (B-6) HRP | Santa Cruz Biotechnology, Dallas, TX | sc-7298 HRP |
| Alexa Fluor™ 488 Phalloidin | ThermoFisher Scientific | A12379 |
| Secondary antibody and other reagents | Source | Identifier/catalog number |
| Goat Anti-Mouse IgG (H+L)-HRP Conjugate | Bio-Rad | 172-1011 |
| Anti-rabbit IgG, HRP-linked Antibody | Cell Signaling | 7074 |
| Bio-Rad colorimetric assay | Bio-Rad | 500-0006 |
| cOmplete Protease Inhibitor Cocktail Tablets | Sigma-Aldrich | 11836153001 |
| Halt™ Phosphatase Inhibitor Cocktail | ThermoFisher | 78420 |
| Criterion TGX 4-20% 26 well gel | Bio-Rad | 5671095 |
| Criterion TGX 4-20% 18 well gel | Bio-Rad | 567-1094 |
| Laemmli sample buffer | Bio-Rad | 161-0737 |
| SuperSignal™ West Femto Maximum Sensitivity Substrate | ThermoFisher Scientific | 34096 |
| SuperSignal™ West Pico PLUS Chemiluminescent Substrate | ThermoFisher Scientific | 34578 |
| Invitrogen™ iBlot™ 2 Gel Transfer Device | ThermoFisher Scientific | IB21001 |
| RIPA Buffer | ThermoFisher Scientific | 89901 |
| Precision Plus protein kaleidoscope marker | Bio-Rad, Hercules, CA | 1610375 |
| **Materials for Flow Cytometry** | | |
| Reagent/Material | Source | Identifier/catalog number |
| CFSE (Carboxyfluorescein succinimidyl ester) | Biolegend, San Diego, California | 423801 |
| live/dead DAPI | ThermoFisher Scientific | L23105 |
| PD-1(Clone:EH12.2H7) | Biolegend | 329906 |
| CD4 (Clone:OKT4) | Biolegend | 317438 |
| CD8 (Clone: RPA-T8) | Biolegend | 301046 |
| CD45 ( Clone:H130) | BD Biosciences | 560367 |
| CD11b (M1/70) | Biolegend | 101235 |
| CD15 (Clone: W6D3) | BD Biosciences | 562369 |
| CD66B (Clone:G10F5) | Biolegend | 305116 |
| CD45 ( Clone:H130) | BD Biosciences | 567401 |
| CD127 (Clone:HIL-7R-M21) | BD Biosciences | 565185 |
| CD4 (Clone:RPA-T4) | BD Biosciences | 561840 |
| FOXP3 (Clone:206D) | Biolegend | 320116 |
| TruStain FCX | Biolegend | 422302 |
| Fixation/ Permeabilization Concentrate | E Bioscience, ThermoFisher Scientific | 00-5123-43 |
| Fixation/ Permeabilization Diluent | E Bioscience, ThermoFisher Scientific | 00-5223-56 |
| Permeabilization Buffer | E Bioscience, ThermoFisher Scientific | 00-8333-56 |
| PBS with 1% BSA | Cell Applications, Inc | 060-500 |
| **Antibodies for CODEX and Immuno Histochemistry** | | |
| CD5-BX017 | Akoya Biosciences, Marlborough, MA | 4250007 |
| CD45-BX007 | Akoya Biosciences | 4150002 |
| CD4-BX026 | Akoya Biosciences | 4250016 |
| CD169-BX015 | Akoya Biosciences | 4350005 |
| CD38-BX019 | Akoya Biosciences | 4150013 |
| CD8a-BX029 | Akoya Biosciences | 4250017 |
| CD11c-BX030 | Akoya Biosciences | 4350013 |
| CD11b-BX025 | Akoya Biosciences | 4150015 |
| CD31-BX002 | Akoya Biosciences | 4250001 |
| Ly6g-BX024 | Akoya Biosciences | 4350015 |
| NOS2 | Akoya Biosciences | 4250073 |
| CD44 | Akoya Biosciences | 4250002 |
| CD206 | ThermoFisher Scientific | MA5-16871(clone:MR5D3) |
| CD49f | Akoya Biosciences | 4550102 |
| CD24 | Akoya Biosciences | 4150014 |
| CD19 | Akoya Biosciences | 4550099 |
| CD44v6 | eBioscience | BMS145 (clone: 9A4) |
| Vimentin | Akoya Biosciences | 4450050 |

| | | |
|---|---|---|
| Anti- human MPO antibody | Agilent, Santa Clara, CA | A0398 |
| Bond Polymer Refine Detection kit | Leica Biosystems | DS9800 |
| **Others** | | |
| Equipment | Source | |
| BD LSRFortessa™ Cell Analyzer | BD biosciences, San Jose, CA | |
| BD FACSVerse Flow Cytometer | BD biosciences, San Jose, CA | |
| Odyssey Fc imager | LI-COR, Lincoln, NE | |
| Cellometer K2 | Nexcelom Bioscience, Lawrence, MA | |
| SpectraMax$^{R}$ iD3 microplate reader | Molecular Devices, LLC, San Jose, CA | |
| TCS SP8 Dive Spectral Microscope | Leica | |
| Software | Source | |
| FlowJo | FlowJo, LLC (Ashland, OR) | |
| Image j | National Institutes of Health | |
| Imaris | BITPLANE (Oxford Instruments Company) | |
| LAS X software | Leica, USA | |
| **Materials for Intravital imaging** | | |
| Reagent/Material | Source | Identifier/catalog number |
| CellTracker Red CMTPX | ThermoFisher Scientific | C34552 |
| xylazine (Anased LA) injection | Vet One, West Pasadena Drive<br>Boise, Idaho 83705 | 510004 |
| Saline Solution | Quality Biological, Gaithersburg, MD | 114-055-101 |
